## Supplemental Figures for "Molecular and cellular similarities in the brain of SARS-CoV-2 and Alzheimer’s disease individuals"

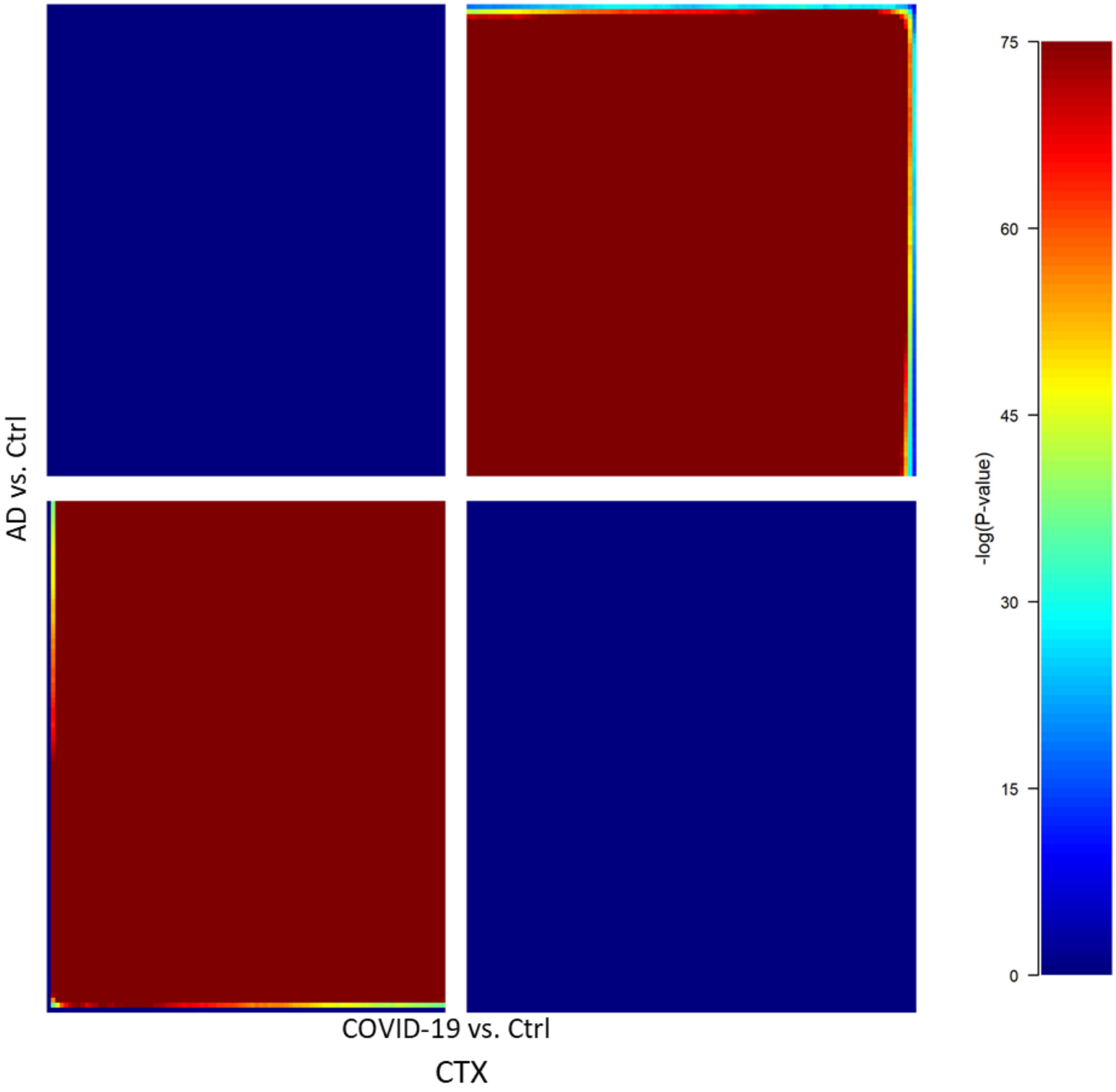


Supplemental Figure 1: RRHO analysis of COVID-19/ Ctrl and AD/ Ctrl, each containing (39,901 DEGs) revealed that gene regulation between the AD/Control and Covid/Control groups showed a positive correlation in CTX region.

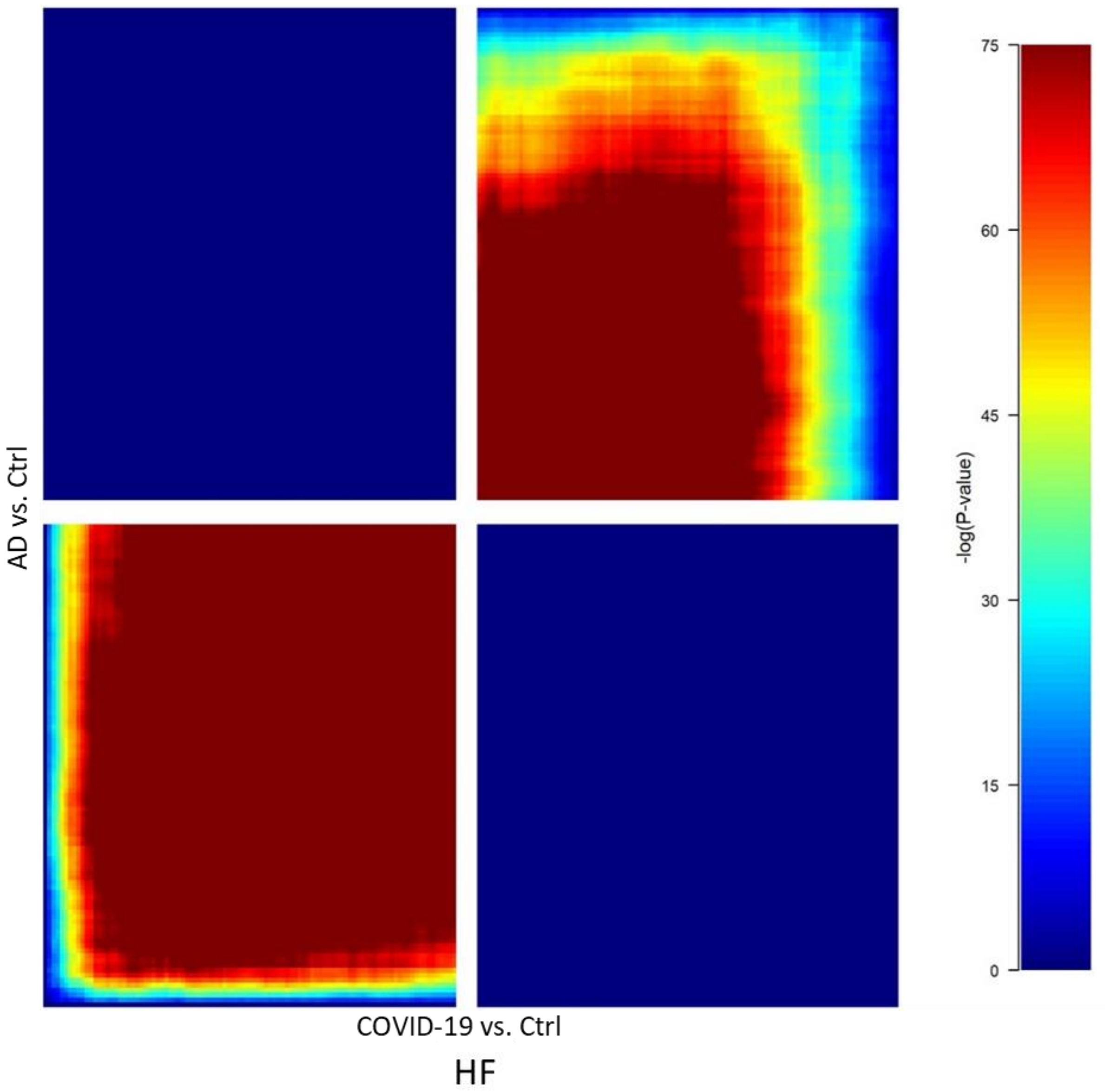


Supplemental Figure 2: RRHO analysis of COVID-19/ Ctrl and AD/ Ctrl, each containing (39,901 DEGs) revealed that gene regulation between the AD/Control and Covid/Control groups showed a positive correlation in HF region.
