## Supplemental Table 1 for "Molecular and cellular similarities in the brain of SARS-CoV-2 and Alzheimer’s disease individuals"

Supplemental Table 1: Demographics of Postmortem Cases

| Characteristic/test | AD | AD+ COVID-19 | COVID-19 | Control |
|---|---|---|---|---|
| Age, years | 82.3 ± 13.3 | 81.3 ± 10.1 | 78.6 ± 11.3 | 76 ± 4.5 |
| Female sex, percent | 50 | 50 | 50 | 50 |
| PMI, hours | 26 ± 23.6 | 29 ± 28 | 33.8 ± 35.6 | 15.7 ± 6.7 |
| ABC | | | | |
| Amyloid | 2.8 ± 0.5 | 2 ± 1.4 | N/a | N/a |
| Braak and Brook | 2.5 ± 0.6 | 2 ± 1.4 | N/a | N/a |
| CERAD | 1.8 ± 1.6 | 2 ± 1.4 | N/a | N/a |
| COVID-19 OtD, days | N/a | 32 ± 19.5 | 27 ± 19.2 | N/a |

Numbers presented are the average per group and standard deviation PMI is post mortem interval (n=4); OtD is onset to death. ABC is a multipoint measurement where A is a measure of amyloid beta deposition, B is a measure of neurofibrillary degeneration based on the Braak score, and C is scored based on neuritic plaques outlined by the Consortium to Establish a Registry for Alzheimer's Disease diagnosis (CERAD).
