## Supplemental Table 2 for "Molecular and cellular similarities in the brain of SARS-CoV-2 and Alzheimer’s disease individuals"

Supplemental Table 2: Blood Chemistry and Symptoms

| | Control | AD | COVID-19 | AD+COVID-19 |
|---|---|---|---|---|
| Symptoms | | | | |
| Fever/ Chills | N/a | N/a | 100 | 75 |
| Ache/ Myalgia | N/a | N/a | 50 | 25 |
| Shortness of Breath | N/a | N/a | 100 | 75 |
| Encephalopathy/ Confusion | N/a | N/a | 100 | 100 |
| Impaired responsiveness/ Coma | N/a | N/a | 100 | 75 |
| Agitation | N/a | N/a | 0 | 100 |
| Anosmia | N/a | N/a | 25 | 0 |
| Chemistry | | | | |
| **D-dimer** | **N/a** | **N/a** | **10.6 ± 6.4** | **12.9 ± 8.9** |
| Ferritin | N/a | N/a | 1887.8 ± 1186.5 | 1713.7 ± 2094.7 |
| Proocalcitonin | N/a | N/a | 12.4 ± 14.5 | 5.7 ± 7.9 |
| ALP | N/a | N/a | 136.3 ± 55.2 | 138.3 ± 106.6 |
| ALT | N/a | N/a | 71.0 ± 27.6 | 122.3 ± 168.0 |
| AST | N/a | N/a | 110.0 ± 38.2 | 116.5 ± 108.8 |
| **CRP** | **N/a** | **N/a** | **364.1 ± 96.7** | **223.4 ± 36.3** |
| Calcium | N/a | N/a | 7.1 ± 0.2 to 9.2 ± 0.5 | 7.7 ± 0.7 to 8.8 ± 0.6 |
| Creatinine | N/a | N/a | 3.6 ± 2.1 | 1.5 ± 0.6 |
| ESR | N/a | N/a | 66.0 ± 46.7 | 69.3 ± 17.0 |
| Eosinophil | N/a | N/a | 0 ± 0 to 0.3 ± 0.4 | 0.0 ± 0.0 to 0.2 ± 0.2 |
| Fibrinogen | N/a | N/a | 0.9 ±0.3 | 0.6 ± 0.0 |
| IL-6 | N/a | N/a | 750.6 ± 1279.7 | 239.4 ± 152.2 |
| INR | N/a | N/a | 1.2 ± 0.2 to 1.9 ± 0.2 | 1.3 ± 0.2 to 1.8 ± 0.5 |
| **LDH** | **N/a** | **N/a** | **748.5 ± 98.0** | **698.0 ± 0.6** |
| Lymphocyte | N/a | N/a | 0.4 ± 0.4 to 1.5 ± 1.2 | 0.8 ± 0.8 to 1.2 ± 0.6 |
| Monocyte | N/a | N/a | 0.1 ± 0.1 to 0.8 ± 0.4 | 0.3 ± 0.3 to 0.8 ± 0.6 |
| **Neutrophil** | **N/a** | **N/a** | **5.0 ± 0.5 to 18.7 ± 9.8** | **5.7 ± 3.5 to 11.2 ± 6.0** |
| Sodium | N/a | N/a | 134.3 ± 7.4 to 154.5 ± 6.7 | 143.0 ± 8.5 to 159.3 ± 10.5 |
| Tropinin | N/a | N/a | 0.8 ± 1.4 | 1.3 ± 2.3 |
| WBC | N/a | N/a | 34.7 ± 16.7 | 13.7 ± 6.4 |
| BUN | N/a | N/a | 106.8 ± 33.7 | 55.7 ± 11.9 |
| Glucose | N/a | N/a | 106.5 ± 33.5 to 304.0 ± 73.3 | 123.3 ± 44.9 to 231.3 ± 60.7 |
| Potassium | N/a | N/a | 3.8 ± 0.5 to 6.2 ± 0.9 | 3.5 ± 0.6 to 4.7 ± 0.1 |
| Albumin | N/a | N/a | 1.5 ± 0.1 | 1.9 ± 0.1 |
| **Hemoglobin** | **N/a** | **N/a** | **8.0 ± 1.6** | **9.0 ± 2.3** |
| Platelets | N/a | N/a | 121.3 ± 78.1 | 273.8 ± 186.8 |

Symptoms are presented as a percentage of the group (n=4); Chemistry numbers are presented as the average per group and standard deviation, based on maximum values collected for single values and the minimum and maximum range collected in cells with two values. Bolded values are implicated as indicators of severe COVID-19 outcomes; ALP is alkaline phosphatase, ALT is alanine aminotransferase, AST is aspartate transferase, CRP is c-reactive protein, ESR is erythrocyte sedimentation, INR international normalized ratio, LDH is lactate dehydrogenase, WBC is white blood count, BUN is blood urea nitrogen.
