## Supplemental Data 1 for "Molecular and cellular similarities in the brain of SARS-CoV-2 and Alzheimer’s disease individuals"

Supplemental Data 1: Gene regulation within the CTX

| | AD+COVID-19 vs. Ctrl | | COVID-19 vs. Ctrl | | AD vs. Ctrl | |
|---|---|---|---|---|---|---|
| Gene | Expr. Log Ratio | Expr. p-value | Expr. Log Ratio | Expr. p-value | Expr. Log Ratio | Expr. p-value |
| ABHD17C | -0.348 | 9.0E-04 | -0.179 | 1.5E-02 | -0.175 | 2.0E-02 |
| ABT1 | -0.564 | 8.0E-03 | -0.293 | 2.7E-02 | -0.278 | 4.5E-02 |
| ACAN | 1.14 | 1.2E-03 | 0.562 | 1.3E-02 | 0.593 | 7.6E-03 |
| ACKR4 | -0.482 | 2.6E-02 | -0.234 | 2.7E-02 | -0.25 | 1.9E-02 |
| ACSS2 | 0.708 | 2.4E-05 | 0.331 | 4.0E-03 | 0.359 | 2.9E-03 |
| ACTL6A | 0.707 | 1.6E-03 | 0.302 | 8.5E-03 | 0.377 | 1.2E-03 |
| ACTR2 | -0.441 | 3.8E-04 | -0.272 | 9.4E-04 | -0.178 | 2.9E-02 |
| ACTR3B | -0.444 | 7.2E-04 | -0.265 | 2.3E-03 | -0.191 | 2.9E-02 |
| ADAM28 | 1.195 | 3.2E-03 | 0.703 | 8.5E-04 | 0.478 | 2.4E-02 |
| ADAT2 | -0.706 | 2.4E-03 | -0.403 | 4.3E-03 | -0.315 | 2.6E-02 |
| ADAT3 | 0.953 | 9.4E-04 | 0.405 | 1.9E-02 | 0.557 | 1.3E-03 |
| ADIPOR2 | 0.776 | 2.3E-03 | 0.414 | 6.9E-03 | 0.357 | 2.3E-02 |
| ADRB2 | 0.783 | 2.4E-03 | 0.369 | 4.4E-02 | 0.359 | 4.5E-02 |
| AGO2 | 0.367 | 1.2E-03 | 0.158 | 2.9E-02 | 0.198 | 6.3E-03 |
| AHNAK | 0.792 | 1.6E-04 | 0.464 | 3.3E-03 | 0.329 | 3.8E-02 |
| AKAP3 | 1.566 | 6.4E-06 | 0.878 | 1.1E-04 | 0.599 | 8.7E-03 |
| AKNA | 0.498 | 9.4E-04 | 0.243 | 3.6E-02 | 0.237 | 3.9E-02 |
| ALKAL2 | 1.114 | 5.4E-04 | 0.432 | 4.9E-02 | 0.617 | 3.3E-03 |
| ANAPC16 | 0.632 | 1.5E-03 | 0.364 | 2.6E-03 | 0.253 | 3.7E-02 |
| ANKRD1 | -2.112 | 1.9E-03 | -0.822 | 4.1E-02 | -0.922 | 2.2E-02 |
| ANKRD20A4P (ir | -0.924 | 3.0E-02 | -0.449 | 4.5E-02 | -0.677 | 3.0E-02 |
| ANKZF1 | 0.601 | 2.1E-03 | 0.331 | 7.9E-03 | 0.261 | 3.7E-02 |
| ANLN | 1.658 | 1.5E-03 | 1.046 | 4.4E-04 | 0.66 | 3.4E-02 |
| ANO10 | -0.27 | 1.8E-03 | -0.157 | 2.4E-03 | -0.122 | 2.1E-02 |
| AP4B1 | -0.5 | 3.9E-03 | -0.211 | 4.3E-02 | -0.291 | 5.4E-03 |
| APOD | 0.693 | 2.5E-02 | 0.378 | 2.7E-02 | 0.353 | 4.1E-02 |
| APOO | -0.344 | 3.2E-03 | -0.221 | 3.9E-04 | -0.135 | 2.9E-02 |
| ARAP1 | 0.827 | 2.4E-06 | 0.44 | 2.7E-06 | 0.372 | 6.9E-05 |
| ARAP1-AS2 | 0.935 | 1.3E-04 | 0.594 | 1.0E-05 | 0.315 | 2.0E-02 |
| ARF3 | -0.633 | 1.6E-04 | -0.295 | 2.1E-02 | -0.349 | 5.0E-03 |
| ARHGAP15 | 0.852 | 2.1E-04 | 0.51 | 2.0E-03 | 0.333 | 4.5E-02 |
| ARHGAP17 | 0.613 | 1.8E-06 | 0.266 | 6.4E-04 | 0.339 | 3.2E-05 |
| ARHGAP21 | 0.437 | 4.3E-03 | 0.202 | 2.4E-02 | 0.224 | 1.6E-02 |
| ARHGAP36 | -2.01 | 7.6E-05 | -1.129 | 1.1E-02 | -1.064 | 1.7E-02 |
| ARHGAP45 | 1.037 | 3.3E-06 | 0.615 | 2.5E-04 | 0.408 | 1.5E-02 |
| ARL1 | -0.477 | 2.2E-04 | -0.263 | 2.0E-03 | -0.224 | 9.7E-03 |
| ARL6 | -0.583 | 2.2E-03 | -0.319 | 1.2E-02 | -0.279 | 2.4E-02 |
| ARL6IP1 | -0.52 | 8.4E-04 | -0.284 | 8.0E-03 | -0.236 | 2.5E-02 |
| ARMC8 | 0.417 | 2.4E-04 | 0.2 | 9.2E-04 | 0.209 | 2.8E-03 |
| ARPC1B | 0.822 | 3.1E-04 | 0.441 | 1.4E-02 | 0.356 | 4.6E-02 |
| ARRDC2 | 0.907 | 4.8E-03 | 0.395 | 4.7E-02 | 0.559 | 4.8E-03 |
| ARSG | -1.01 | 1.4E-05 | -0.555 | 1.2E-03 | -0.493 | 4.2E-03 |
| ATAD1 | -0.369 | 7.9E-05 | -0.227 | 6.2E-05 | -0.151 | 8.0E-03 |
| ATF7 | 0.511 | 5.6E-03 | 0.265 | 1.4E-02 | 0.236 | 3.0E-02 |
| ATF7-NPFF | 0.518 | 5.0E-03 | 0.272 | 1.2E-02 | 0.237 | 3.0E-02 |
| ATG4C | 0.795 | 4.4E-03 | 0.42 | 1.6E-02 | 0.362 | 4.6E-02 |
| ATOH8 | 1.025 | 9.9E-04 | 0.54 | 1.2E-02 | 0.548 | 1.1E-02 |

| | | | | | | |
|---|---|---|---|---|---|---|
| ATP11A | 0.675 | 1.4E-04 | 0.413 | 2.6E-04 | 0.249 | 2.8E-02 |
| ATP2B1-AS1 | -0.995 | 1.1E-04 | -0.47 | 1.0E-02 | -0.563 | 1.5E-03 |
| ATP6AP1 | -0.495 | 1.4E-02 | -0.263 | 2.1E-02 | -0.242 | 3.0E-02 |
| ATP6AP2 | -0.589 | 4.4E-05 | -0.35 | 4.4E-04 | -0.243 | 1.3E-02 |
| ATP6V0E2-AS1 | -0.401 | 1.0E-04 | -0.219 | 8.0E-03 | -0.199 | 1.7E-02 |
| ATP6V1E1 | -0.49 | 7.8E-04 | -0.238 | 3.4E-03 | -0.264 | 9.1E-04 |
| ATRN | -0.326 | 2.7E-03 | -0.153 | 9.1E-03 | -0.184 | 1.1E-03 |
| AUNIP | -0.378 | 3.2E-03 | -0.189 | 4.3E-02 | -0.203 | 2.7E-02 |
| AZGP1 | 1.797 | 8.7E-03 | 0.947 | 2.2E-02 | 0.923 | 2.6E-02 |
| B3GALNT1 | -0.674 | 1.3E-04 | -0.285 | 1.9E-02 | -0.398 | 9.3E-04 |
| B3GAT1-DT | -0.939 | 3.3E-03 | -0.553 | 2.9E-03 | -0.5 | 6.9E-03 |
| B4GALT6 | -0.699 | 1.7E-03 | -0.371 | 1.2E-02 | -0.325 | 2.7E-02 |
| BABAM2 | -0.341 | 5.3E-04 | -0.161 | 2.8E-03 | -0.192 | 2.5E-04 |
| BAHCC1 | 0.628 | 4.8E-04 | 0.395 | 3.0E-05 | 0.228 | 1.5E-02 |
| BANP | 0.871 | 1.4E-03 | 0.384 | 3.8E-02 | 0.416 | 2.5E-02 |
| BARD1 | 0.921 | 3.0E-06 | 0.337 | 4.9E-02 | 0.556 | 1.1E-03 |
| BEND6 | -0.523 | 4.7E-08 | -0.31 | 5.9E-03 | -0.224 | 4.7E-02 |
| BEX4 | -0.564 | 1.7E-04 | -0.292 | 2.2E-03 | -0.278 | 2.9E-03 |
| BEX5 | -1.117 | 8.8E-04 | -0.704 | 1.6E-03 | -0.467 | 3.9E-02 |
| BICRA | 0.367 | 7.7E-04 | 0.236 | 2.4E-05 | 0.121 | 3.1E-02 |
| BMP1 | 0.658 | 6.6E-03 | 0.32 | 2.0E-02 | 0.318 | 2.1E-02 |
| BOK | 1.463 | 5.5E-04 | 0.926 | 2.3E-05 | 0.524 | 1.7E-02 |
| BPHL | -0.57 | 1.9E-02 | -0.292 | 4.9E-02 | -0.314 | 3.4E-02 |
| BRD9 | 0.463 | 1.7E-02 | 0.257 | 4.3E-03 | 0.181 | 4.2E-02 |
| BRF1 | 0.388 | 5.0E-05 | 0.188 | 1.1E-03 | 0.19 | 1.0E-03 |
| BSCL2 | -0.693 | 5.1E-04 | -0.374 | 1.9E-02 | -0.341 | 3.2E-02 |
| BTC | -0.97 | 2.3E-05 | -0.622 | 7.1E-04 | -0.377 | 4.2E-02 |
| C1QTNF2 | -0.881 | 1.1E-04 | -0.401 | 2.2E-02 | -0.527 | 2.3E-03 |
| CA2 | 1.058 | 1.5E-03 | 0.406 | 3.2E-02 | 0.641 | 2.1E-03 |
| CAPN15 | 0.667 | 5.8E-03 | 0.352 | 1.1E-02 | 0.313 | 2.1E-02 |
| CARD9 | 1.222 | 1.5E-05 | 0.494 | 2.8E-02 | 0.716 | 1.8E-03 |
| CARHSP1 | 0.599 | 2.2E-04 | 0.316 | 3.3E-04 | 0.279 | 1.5E-03 |
| CASC3 | 0.453 | 1.0E-04 | 0.255 | 6.5E-04 | 0.188 | 1.1E-02 |
| CASP2 | 0.339 | 5.4E-04 | 0.143 | 2.6E-02 | 0.183 | 6.8E-03 |
| CCDC113 | -0.638 | 3.6E-04 | -0.322 | 3.9E-02 | -0.324 | 3.4E-02 |
| CCDC32 | -0.491 | 4.3E-04 | -0.312 | 7.6E-05 | -0.189 | 1.7E-02 |
| CCPG1 | -0.358 | 2.2E-03 | -0.168 | 7.3E-03 | -0.209 | 1.1E-03 |
| CD226 | -0.553 | 6.6E-04 | -0.281 | 3.6E-02 | -0.291 | 2.8E-02 |
| CD28 | -3.293 | 1.7E-03 | -2.453 | 1.6E-04 | -1.137 | 4.9E-02 |
| CD86 | 1.334 | 1.0E-03 | 0.745 | 1.6E-03 | 0.655 | 5.0E-03 |
| CDC5L | -0.348 | 1.7E-05 | -0.228 | 1.9E-05 | -0.134 | 9.0E-03 |
| CDK19 | 0.65 | 2.7E-03 | 0.359 | 7.5E-03 | 0.28 | 4.5E-02 |
| CDK5RAP2 | 0.436 | 1.9E-03 | 0.18 | 1.6E-02 | 0.248 | 9.7E-04 |
| CDK6-AS1 | -1.135 | 5.5E-03 | -0.608 | 1.5E-02 | -0.551 | 2.7E-02 |
| CEACAM8 | -1.201 | 1.8E-03 | -0.561 | 1.1E-02 | -0.587 | 7.7E-03 |
| CEACAMP5 | -1.039 | 5.5E-04 | -0.518 | 2.6E-03 | -0.54 | 2.1E-03 |
| CEP104 | 0.361 | 1.7E-04 | 0.165 | 1.1E-03 | 0.19 | 2.3E-04 |
| CEP152 | 0.738 | 2.4E-03 | 0.298 | 4.0E-02 | 0.428 | 3.1E-03 |
| CEP192-DT | -0.591 | 1.2E-04 | -0.339 | 2.5E-03 | -0.269 | 1.4E-02 |
| CEP295NL | 0.761 | 7.5E-05 | 0.375 | 9.0E-03 | 0.382 | 6.9E-03 |

| | | | | | | |
|---|---|---|---|---|---|---|
| CEP43 | -0.525 | 1.1E-04 | -0.274 | 9.1E-04 | -0.257 | 1.4E-03 |
| CEP68 | -0.274 | 2.9E-03 | -0.135 | 1.8E-02 | -0.152 | 8.5E-03 |
| CERNA2 | 3.295 | 5.0E-05 | 1.161 | 1.6E-02 | 1.456 | 3.8E-03 |
| CFL1P1 | -0.582 | 2.3E-03 | -0.343 | 4.2E-03 | -0.245 | 4.1E-02 |
| CHD7 | 0.796 | 4.6E-03 | 0.429 | 8.8E-03 | 0.359 | 2.8E-02 |
| CHDH | 0.92 | 7.7E-06 | 0.398 | 6.9E-03 | 0.5 | 5.8E-04 |
| CHEK2 | 1.021 | 2.4E-04 | 0.43 | 1.1E-02 | 0.573 | 8.0E-04 |
| CHFR | 0.288 | 2.4E-03 | 0.147 | 1.3E-02 | 0.131 | 2.5E-02 |
| CHRM5 | 1.785 | 6.0E-04 | 1.027 | 4.8E-03 | 0.849 | 2.1E-02 |
| CHRNA7 | -1.897 | 7.6E-05 | -1.313 | 3.4E-05 | -0.677 | 3.4E-02 |
| CHST3 | 1.244 | 1.3E-04 | 0.675 | 4.4E-03 | 0.571 | 1.6E-02 |
| CLASRP | 0.727 | 5.1E-11 | 0.341 | 2.2E-04 | 0.375 | 2.6E-05 |
| CLCN2 | 0.524 | 1.4E-03 | 0.225 | 4.6E-02 | 0.285 | 1.2E-02 |
| CLCN7 | 0.453 | 5.2E-03 | 0.204 | 4.9E-02 | 0.241 | 2.1E-02 |
| CLDN15 | 1.284 | 1.7E-05 | 0.741 | 4.4E-06 | 0.522 | 1.1E-03 |
| CLIP2 | 0.601 | 2.5E-05 | 0.309 | 7.1E-04 | 0.282 | 1.3E-03 |
| CLK1 | 0.669 | 1.8E-04 | 0.286 | 4.0E-03 | 0.365 | 2.4E-04 |
| CLMN | 1.206 | 1.2E-03 | 0.584 | 9.3E-03 | 0.618 | 6.8E-03 |
| CMTM2 | -1.154 | 6.2E-04 | -0.622 | 2.3E-03 | -0.572 | 6.1E-03 |
| COA7 | -0.634 | 1.1E-03 | -0.325 | 7.4E-03 | -0.322 | 6.9E-03 |
| COPG1 | -0.387 | 2.5E-03 | -0.224 | 1.2E-02 | -0.179 | 4.0E-02 |
| COPS7A | -0.677 | 4.0E-07 | -0.313 | 2.1E-04 | -0.377 | 4.1E-06 |
| COPS8 | -0.444 | 1.2E-03 | -0.236 | 8.2E-03 | -0.217 | 1.2E-02 |
| COX7B | -0.636 | 2.5E-04 | -0.337 | 1.2E-03 | -0.315 | 2.3E-03 |
| CPM | 0.891 | 8.1E-03 | 0.4 | 4.2E-02 | 0.509 | 9.8E-03 |
| CPQ | 0.631 | 8.4E-04 | 0.334 | 1.7E-03 | 0.287 | 8.3E-03 |
| CPT1B | 0.826 | 5.1E-04 | 0.347 | 1.8E-02 | 0.476 | 1.2E-03 |
| CPZ | -1.383 | 3.8E-03 | -0.703 | 1.8E-02 | -0.655 | 2.8E-02 |
| CSRNP2 | -0.316 | 5.2E-05 | -0.177 | 1.1E-02 | -0.153 | 2.8E-02 |
| CSRP1 | 0.878 | 4.6E-05 | 0.367 | 2.4E-02 | 0.503 | 2.0E-03 |
| CTBP2 | 0.601 | 5.8E-06 | 0.252 | 3.4E-03 | 0.337 | 9.1E-05 |
| CTDSP2 | 0.558 | 4.6E-04 | 0.357 | 1.0E-05 | 0.191 | 1.7E-02 |
| CTDSPL2 | 0.276 | 2.1E-03 | 0.139 | 1.6E-03 | 0.126 | 8.7E-03 |
| CTTN | 0.417 | 8.5E-05 | 0.192 | 7.9E-03 | 0.212 | 3.2E-03 |
| CUL2 | -0.202 | 1.7E-02 | -0.111 | 3.8E-02 | -0.101 | 4.6E-02 |
| DAZAP1 | 0.547 | 4.2E-06 | 0.325 | 8.3E-05 | 0.209 | 1.1E-02 |
| DCAF4 | -0.614 | 2.2E-04 | -0.321 | 2.0E-03 | -0.3 | 4.5E-03 |
| DDB2 | 0.789 | 7.2E-04 | 0.385 | 3.9E-02 | 0.374 | 4.4E-02 |
| DDX19A-DT | 0.457 | 2.4E-03 | 0.234 | 1.1E-02 | 0.208 | 2.5E-02 |
| DDX39A | 0.594 | 1.1E-03 | 0.277 | 2.6E-02 | 0.309 | 1.3E-02 |
| DEK | 0.499 | 8.5E-04 | 0.22 | 9.2E-03 | 0.265 | 1.8E-03 |
| DIO1 | 0.929 | 9.3E-05 | 0.431 | 6.3E-03 | 0.454 | 3.9E-03 |
| DIP2C | 0.558 | 2.9E-05 | 0.271 | 4.0E-04 | 0.277 | 4.3E-04 |
| DIXDC1 | 0.493 | 4.3E-03 | 0.212 | 3.6E-02 | 0.267 | 1.2E-02 |
| DLAT | -0.245 | 4.7E-03 | -0.151 | 4.2E-03 | -0.103 | 4.9E-02 |
| DLD | -0.308 | 3.5E-03 | -0.176 | 4.3E-03 | -0.142 | 2.1E-02 |
| DLX1 | -1.205 | 3.6E-04 | -0.661 | 1.1E-02 | -0.58 | 2.7E-02 |
| DNAAF4-CCPG1 | -0.374 | 3.3E-04 | -0.175 | 3.6E-03 | -0.218 | 3.2E-04 |
| DNAJA2 | -0.459 | 3.0E-03 | -0.274 | 3.5E-03 | -0.191 | 3.4E-02 |
| DNAJB14 | -0.298 | 2.5E-03 | -0.161 | 3.0E-02 | -0.145 | 4.4E-02 |

| | | | | | | |
|---|---|---|---|---|---|---|
| DNAJC18 | -0.385 | 1.9E-03 | -0.18 | 1.2E-02 | -0.216 | 2.8E-03 |
| DNAJC24 | -0.424 | 4.5E-03 | -0.193 | 3.7E-02 | -0.243 | 1.3E-02 |
| DNM2 | 0.502 | 2.7E-06 | 0.26 | 4.7E-04 | 0.231 | 1.5E-03 |
| DOCK11 | 0.647 | 1.1E-05 | 0.272 | 4.2E-02 | 0.376 | 4.9E-03 |
| DOT1L | 0.62 | 1.3E-04 | 0.258 | 2.5E-02 | 0.353 | 2.0E-03 |
| DPY19L2P3 | -0.526 | 1.4E-05 | -0.274 | 6.3E-03 | -0.264 | 8.6E-03 |
| DTWD2 | -0.545 | 2.4E-03 | -0.322 | 4.1E-04 | -0.245 | 6.9E-03 |
| DUSP22 | 0.4 | 2.4E-03 | 0.201 | 2.1E-02 | 0.189 | 3.0E-02 |
| DVL2 | 0.458 | 7.5E-04 | 0.197 | 1.2E-02 | 0.251 | 7.9E-04 |
| DYNC2LI1 | -0.53 | 8.4E-04 | -0.27 | 1.3E-03 | -0.288 | 5.1E-04 |
| E2F3 | -0.636 | 1.7E-04 | -0.319 | 8.7E-03 | -0.332 | 6.5E-03 |
| EBF4 | 0.485 | 1.3E-02 | 0.244 | 2.8E-02 | 0.219 | 4.4E-02 |
| EBLN2 | 0.984 | 6.3E-04 | 0.566 | 2.0E-03 | 0.386 | 4.0E-02 |
| EBLN3P | -0.333 | 3.5E-04 | -0.181 | 3.0E-03 | -0.167 | 3.6E-03 |
| EDA2R | 1.148 | 3.4E-04 | 0.505 | 4.0E-02 | 0.556 | 2.8E-02 |
| EHBP1L1 | 0.529 | 1.9E-03 | 0.28 | 2.2E-02 | 0.24 | 4.2E-02 |
| EHMT1 | 0.332 | 1.0E-03 | 0.194 | 3.2E-04 | 0.125 | 2.1E-02 |
| EIF3M | -0.314 | 6.5E-03 | -0.192 | 3.6E-03 | -0.133 | 4.3E-02 |
| EIF4E3 | -0.331 | 6.4E-03 | -0.161 | 3.3E-02 | -0.179 | 1.8E-02 |
| ELF1 | 0.795 | 6.0E-04 | 0.373 | 5.6E-03 | 0.408 | 2.4E-03 |
| ELF2 | 0.422 | 8.3E-04 | 0.211 | 2.0E-02 | 0.195 | 3.2E-02 |
| ELMOD3 | 0.525 | 1.8E-05 | 0.266 | 4.6E-04 | 0.25 | 1.0E-03 |
| ELOA-AS1 | 0.596 | 7.3E-05 | 0.203 | 2.4E-02 | 0.368 | 3.1E-05 |
| EML2 | 0.465 | 4.3E-03 | 0.206 | 3.4E-02 | 0.244 | 1.2E-02 |
| ENPP5 | -0.58 | 3.5E-03 | -0.349 | 3.3E-03 | -0.234 | 4.7E-02 |
| EPHA5-AS1 | -0.889 | 6.4E-04 | -0.478 | 2.2E-02 | -0.479 | 2.1E-02 |
| ERBIN | 1.039 | 1.1E-04 | 0.446 | 1.3E-02 | 0.59 | 1.1E-03 |
| ERFE | -1.055 | 3.2E-03 | -0.562 | 1.3E-02 | -0.555 | 1.6E-02 |
| ERI3 | 0.295 | 1.2E-02 | 0.146 | 2.5E-02 | 0.139 | 3.5E-02 |
| ERICH3-AS1 | -1.081 | 1.7E-05 | -0.525 | 3.7E-02 | -0.613 | 1.3E-02 |
| ERICH6B | -0.462 | 1.6E-02 | -0.238 | 4.8E-02 | -0.244 | 4.2E-02 |
| ETV5-AS1 | -0.776 | 3.9E-03 | -0.349 | 4.5E-02 | -0.455 | 8.9E-03 |
| EVI5L | 0.433 | 1.1E-02 | 0.259 | 3.3E-03 | 0.174 | 3.7E-02 |
| EXT1 | -0.436 | 1.2E-05 | -0.229 | 2.2E-02 | -0.222 | 2.7E-02 |
| FAM133B | 0.495 | 4.0E-03 | 0.224 | 1.6E-02 | 0.251 | 6.4E-03 |
| FAM170B-AS1 | 2.291 | 2.9E-04 | 0.992 | 6.4E-03 | 1.324 | 2.8E-04 |
| FAM174A | -0.416 | 2.5E-03 | -0.225 | 2.0E-02 | -0.202 | 3.8E-02 |
| FAM229A | 1.066 | 9.5E-05 | 0.611 | 3.2E-04 | 0.493 | 3.8E-03 |
| FAM234B | -0.701 | 5.8E-04 | -0.346 | 3.1E-02 | -0.374 | 1.9E-02 |
| FAM3D-AS1 | 1.255 | 4.6E-04 | 0.562 | 1.3E-02 | 0.697 | 2.1E-03 |
| FAM53B | 0.724 | 2.4E-04 | 0.387 | 6.2E-04 | 0.33 | 3.6E-03 |
| FAM53B-AS1 | 0.653 | 3.5E-03 | 0.37 | 7.9E-03 | 0.293 | 3.8E-02 |
| FANCB | 0.964 | 1.2E-04 | 0.427 | 2.4E-03 | 0.547 | 1.6E-04 |
| FANCE | 0.659 | 5.5E-03 | 0.349 | 1.3E-02 | 0.306 | 3.0E-02 |
| FASTKD2 | -0.397 | 5.8E-04 | -0.22 | 7.9E-03 | -0.189 | 2.4E-02 |
| FBRSL1 | 0.558 | 1.3E-03 | 0.29 | 8.9E-03 | 0.255 | 2.3E-02 |
| FBXL21P | -1.863 | 1.2E-03 | -1.078 | 9.2E-03 | -1.086 | 8.0E-03 |
| FCHO1 | 0.906 | 6.8E-04 | 0.455 | 3.1E-03 | 0.439 | 4.3E-03 |
| FFAR4 | -0.963 | 2.5E-04 | -0.531 | 1.6E-02 | -0.471 | 3.4E-02 |
| FGD3 | 0.943 | 3.6E-04 | 0.445 | 6.8E-03 | 0.456 | 5.0E-03 |

| | | | | | | |
|---|---|---|---|---|---|---|
| FLACC1 | -0.77 | 1.6E-03 | -0.428 | 5.1E-03 | -0.365 | 1.6E-02 |
| FLII | 0.573 | 1.9E-05 | 0.269 | 1.1E-03 | 0.296 | 3.5E-04 |
| FLJ40194 | 0.997 | 5.2E-04 | 0.479 | 8.9E-03 | 0.525 | 4.6E-03 |
| FNDC5 | -0.602 | 1.3E-04 | -0.239 | 4.3E-02 | -0.394 | 5.0E-04 |
| FOXG1-AS1 | -0.536 | 2.9E-04 | -0.274 | 4.5E-02 | -0.279 | 4.1E-02 |
| FOXO4 | 0.957 | 1.5E-03 | 0.561 | 3.2E-03 | 0.406 | 3.1E-02 |
| FOXP4 | 0.475 | 1.0E-02 | 0.243 | 9.0E-03 | 0.213 | 2.0E-02 |
| FREM3 | -1.255 | 9.7E-08 | -0.831 | 6.6E-06 | -0.441 | 1.7E-02 |
| FRG1JP | 1.906 | 8.8E-05 | 1.133 | 7.6E-04 | 0.705 | 3.6E-02 |
| FRMD6 | -0.693 | 8.7E-07 | -0.391 | 2.9E-03 | -0.321 | 1.3E-02 |
| FRMD6-AS2 | -1.645 | 2.2E-05 | -1.001 | 2.3E-04 | -0.618 | 2.6E-02 |
| GALNT11 | -0.595 | 1.8E-07 | -0.265 | 1.1E-03 | -0.341 | 3.8E-05 |
| GAP43 | -0.832 | 3.3E-08 | -0.424 | 2.3E-02 | -0.414 | 2.7E-02 |
| GAS6 | 0.4 | 7.7E-04 | 0.186 | 2.1E-02 | 0.203 | 1.2E-02 |
| GAS6-AS1 | 0.497 | 3.1E-03 | 0.211 | 4.9E-02 | 0.267 | 1.3E-02 |
| GIPR | 1.461 | 1.3E-03 | 0.623 | 3.7E-02 | 0.802 | 7.7E-03 |
| GNG3 | -1.067 | 3.0E-05 | -0.519 | 3.0E-02 | -0.584 | 1.4E-02 |
| GNPTAB | -0.296 | 6.4E-03 | -0.137 | 4.8E-02 | -0.172 | 1.3E-02 |
| GNRH1 | 1.586 | 1.7E-05 | 0.956 | 2.1E-05 | 0.645 | 5.5E-03 |
| GOLGA3 | 0.334 | 8.9E-03 | 0.167 | 1.6E-02 | 0.157 | 2.3E-02 |
| GOLT1B | -0.59 | 1.8E-03 | -0.39 | 2.5E-04 | -0.219 | 4.0E-02 |
| GOPC | -0.273 | 1.6E-03 | -0.122 | 2.1E-02 | -0.161 | 3.0E-03 |
| GPC6 | -1 | 2.0E-04 | -0.441 | 3.0E-02 | -0.596 | 3.1E-03 |
| GPIHBP1 | 1.603 | 7.3E-03 | 0.805 | 4.3E-02 | 0.929 | 2.1E-02 |
| GPR108 | 0.577 | 3.7E-03 | 0.328 | 8.3E-03 | 0.265 | 3.3E-02 |
| GPR141 | 1.508 | 8.9E-04 | 0.779 | 1.8E-02 | 0.668 | 4.5E-02 |
| GPRASP1 | -0.664 | 1.4E-03 | -0.359 | 2.2E-02 | -0.323 | 3.9E-02 |
| GPRC5B | 0.836 | 5.3E-04 | 0.452 | 1.7E-03 | 0.377 | 8.7E-03 |
| GPSM1 | 0.461 | 2.5E-03 | 0.191 | 4.6E-02 | 0.265 | 8.2E-03 |
| GTF2A1 | -0.387 | 6.8E-04 | -0.22 | 2.0E-04 | -0.173 | 3.1E-03 |
| H1-9P | 1.522 | 6.4E-04 | 0.841 | 2.4E-03 | 0.63 | 2.3E-02 |
| H2AZ1-DT | -0.415 | 1.0E-02 | -0.202 | 4.8E-02 | -0.218 | 3.6E-02 |
| HAPLN1 | -1.29 | 3.1E-03 | -0.628 | 4.4E-02 | -0.688 | 2.5E-02 |
| HAPLN2 | 1.427 | 1.1E-03 | 0.739 | 2.2E-02 | 0.682 | 3.6E-02 |
| HAR1A | -1.246 | 2.7E-03 | -0.633 | 2.6E-02 | -0.582 | 4.1E-02 |
| HAUS5 | 0.74 | 4.8E-03 | 0.367 | 1.3E-02 | 0.341 | 1.9E-02 |
| HBP1 | 0.418 | 1.3E-03 | 0.2 | 8.8E-03 | 0.208 | 6.5E-03 |
| HCLS1 | 1.304 | 4.3E-05 | 0.574 | 6.4E-03 | 0.642 | 2.4E-03 |
| HDAC4 | 0.49 | 3.1E-07 | 0.279 | 1.5E-06 | 0.196 | 7.8E-04 |
| HDGF | 0.401 | 3.7E-03 | 0.178 | 2.7E-02 | 0.216 | 8.3E-03 |
| HEBP2 | 0.501 | 1.4E-03 | 0.269 | 1.5E-02 | 0.221 | 4.4E-02 |
| HEXD | 0.829 | 1.2E-04 | 0.298 | 1.3E-02 | 0.524 | 1.9E-05 |
| HIGD1C | 1.011 | 3.1E-03 | 0.493 | 3.8E-02 | 0.499 | 3.5E-02 |
| HIP1 | 0.967 | 1.5E-03 | 0.604 | 4.6E-04 | 0.359 | 3.7E-02 |
| HMBOX1 | 0.362 | 3.2E-03 | 0.156 | 3.1E-02 | 0.194 | 7.8E-03 |
| HMG20A | -0.291 | 2.4E-03 | -0.15 | 1.8E-02 | -0.154 | 2.1E-02 |
| HMG20B | 0.876 | 1.4E-03 | 0.406 | 9.6E-03 | 0.459 | 3.4E-03 |
| HNRNPA2B1 | 0.366 | 1.7E-04 | 0.14 | 1.5E-02 | 0.212 | 2.3E-04 |
| HNRNPF | 0.605 | 7.7E-05 | 0.218 | 4.9E-02 | 0.378 | 5.8E-04 |
| HNRNPH1 | 0.35 | 5.6E-04 | 0.169 | 2.4E-03 | 0.169 | 5.7E-03 |

| | | | | | | |
|---|---|---|---|---|---|---|
| HNRNPUL2-BSC | -0.492 | 1.4E-03 | -0.255 | 2.3E-02 | -0.257 | 2.1E-02 |
| HPN | 1.53 | 2.9E-04 | 0.806 | 1.6E-02 | 0.691 | 4.1E-02 |
| HTATIP2 | 1.112 | 6.8E-05 | 0.551 | 1.1E-03 | 0.519 | 2.5E-03 |
| ICA1 | -0.726 | 7.2E-04 | -0.363 | 2.2E-02 | -0.411 | 9.3E-03 |
| ICA1L | -0.444 | 5.3E-05 | -0.197 | 3.4E-02 | -0.253 | 6.1E-03 |
| IKZF2 | 0.799 | 1.8E-03 | 0.373 | 1.9E-02 | 0.415 | 1.1E-02 |
| IL10RA | 1.019 | 3.4E-04 | 0.459 | 3.0E-02 | 0.579 | 7.8E-03 |
| INPPL1 | 0.625 | 6.7E-04 | 0.34 | 1.2E-02 | 0.289 | 3.5E-02 |
| INSM1 | -0.99 | 8.7E-03 | -0.454 | 4.6E-02 | -0.539 | 2.2E-02 |
| INSR | 0.508 | 9.2E-05 | 0.264 | 3.0E-04 | 0.234 | 1.2E-03 |
| IP6K3 | 2.099 | 1.2E-05 | 0.937 | 1.3E-03 | 1.155 | 9.2E-05 |
| IRAK2 | 1.115 | 4.6E-05 | 0.62 | 3.6E-03 | 0.473 | 2.6E-02 |
| IRF2 | 0.417 | 9.3E-04 | 0.228 | 3.8E-03 | 0.176 | 2.5E-02 |
| IRF8 | 1.516 | 2.7E-04 | 0.777 | 3.6E-03 | 0.688 | 1.2E-02 |
| ITGA11 | 0.972 | 1.1E-04 | 0.418 | 2.7E-02 | 0.555 | 4.2E-03 |
| ITGB8 | 0.644 | 6.0E-03 | 0.33 | 1.7E-02 | 0.305 | 3.0E-02 |
| ITGB8-AS1 | 0.842 | 2.4E-03 | 0.47 | 7.0E-03 | 0.382 | 2.8E-02 |
| ITPKB | 0.965 | 4.3E-05 | 0.445 | 1.7E-02 | 0.528 | 4.8E-03 |
| ITPKB-IT1 | 0.934 | 1.3E-04 | 0.397 | 4.3E-02 | 0.536 | 6.3E-03 |
| ITPRIP-AS1 | 1.862 | 2.1E-03 | 0.644 | 4.5E-02 | 0.72 | 2.6E-02 |
| IWS1 | 0.526 | 1.7E-03 | 0.268 | 1.6E-03 | 0.246 | 4.1E-03 |
| JAKMIP3 | 0.791 | 8.0E-04 | 0.421 | 4.3E-03 | 0.351 | 1.7E-02 |
| KAT6A | 0.297 | 2.6E-03 | 0.114 | 4.4E-02 | 0.17 | 2.7E-03 |
| KCNJ11 | -0.528 | 1.0E-03 | -0.262 | 4.4E-02 | -0.277 | 3.4E-02 |
| KCNMB4 | 0.939 | 1.2E-03 | 0.446 | 4.8E-02 | 0.486 | 3.4E-02 |
| KCNV2 | -0.726 | 4.6E-03 | -0.47 | 3.3E-04 | -0.291 | 2.5E-02 |
| KDM4B | 0.46 | 3.1E-04 | 0.242 | 1.3E-02 | 0.207 | 2.9E-02 |
| KDM6A | 0.469 | 2.6E-02 | 0.207 | 1.4E-02 | 0.255 | 4.0E-03 |
| KIAA1755 | 1.079 | 2.6E-03 | 0.624 | 4.5E-03 | 0.467 | 3.4E-02 |
| KIAA1958 | 0.56 | 2.0E-03 | 0.245 | 1.1E-02 | 0.303 | 2.3E-03 |
| KIF12 | -1.349 | 3.5E-05 | -0.879 | 1.7E-04 | -0.492 | 4.4E-02 |
| KIF13B | 0.853 | 6.4E-04 | 0.475 | 2.2E-03 | 0.369 | 2.1E-02 |
| KIF1C | 0.836 | 2.5E-03 | 0.497 | 3.0E-04 | 0.331 | 1.4E-02 |
| KIF3B | -0.415 | 4.1E-04 | -0.236 | 2.5E-03 | -0.192 | 1.4E-02 |
| KLF15 | 1.295 | 3.8E-05 | 0.694 | 1.4E-04 | 0.596 | 9.5E-04 |
| KLF8 | -0.647 | 8.1E-04 | -0.311 | 9.6E-03 | -0.353 | 3.4E-03 |
| KLHDC10 | -0.345 | 6.7E-05 | -0.131 | 1.6E-02 | -0.224 | 6.1E-05 |
| KLHL21 | 0.62 | 5.6E-03 | 0.313 | 2.6E-02 | 0.279 | 4.8E-02 |
| KLHL42 | -0.441 | 1.1E-05 | -0.255 | 1.9E-04 | -0.196 | 4.1E-03 |
| KLHL6-AS1 | 3.183 | 7.3E-03 | 1.189 | 2.7E-02 | 1.149 | 3.5E-02 |
| KLK5 | -2.375 | 1.1E-03 | -1.222 | 2.4E-02 | -1.469 | 6.1E-03 |
| KLRC4-KLRK1/K | 1.011 | 2.7E-03 | 0.538 | 4.6E-03 | 0.517 | 4.0E-02 |
| KMT2C | 0.243 | 3.1E-03 | 0.107 | 3.0E-02 | 0.124 | 1.3E-02 |
| L2HGDH | -0.428 | 8.6E-03 | -0.224 | 1.1E-02 | -0.211 | 1.8E-02 |
| LBX2 | 1.638 | 3.2E-03 | 0.734 | 2.0E-02 | 0.921 | 3.6E-03 |
| LCLAT1 | -0.439 | 1.9E-03 | -0.245 | 4.2E-03 | -0.205 | 1.7E-02 |
| LETMD1 | -0.389 | 1.8E-03 | -0.212 | 3.2E-02 | -0.194 | 4.8E-02 |
| LINC00488 | -0.988 | 7.4E-03 | -0.548 | 2.3E-02 | -0.493 | 3.6E-02 |
| LINC00498 | 1.677 | 6.6E-03 | 0.756 | 4.0E-02 | 0.77 | 3.6E-02 |
| LINC00842 | -1.919 | 1.1E-04 | -0.799 | 1.8E-02 | -1.179 | 4.5E-04 |

| | | | | | | |
|---|---|---|---|---|---|---|
| LINC00862 | 1.31 | 2.7E-04 | 0.655 | 2.2E-03 | 0.582 | 6.5E-03 |
| LINC01007 | -1.987 | 4.4E-06 | -0.979 | 1.1E-02 | -1.026 | 8.0E-03 |
| LINC01065 | -1.573 | 9.6E-04 | -0.864 | 5.7E-03 | -0.693 | 2.5E-02 |
| LINC01128 | -0.362 | 2.7E-03 | -0.199 | 2.3E-02 | -0.18 | 4.0E-02 |
| LINC01277 | -0.919 | 2.4E-03 | -0.442 | 2.5E-02 | -0.524 | 8.0E-03 |
| LINC01338 | 1.873 | 4.7E-04 | 1.132 | 2.3E-03 | 0.878 | 2.1E-02 |
| LINC01561 | 1.369 | 2.2E-03 | 0.735 | 2.9E-03 | 0.579 | 1.9E-02 |
| LINC01736 | 2.452 | 7.4E-04 | 1.345 | 5.3E-03 | 1.043 | 3.0E-02 |
| LINC01844 | -1.281 | 4.4E-04 | -0.764 | 4.6E-03 | -0.552 | 4.1E-02 |
| LINC02320 | -1.318 | 8.6E-04 | -0.812 | 3.1E-04 | -0.577 | 9.2E-03 |
| LINC02516 | 1.267 | 1.4E-02 | 0.581 | 3.6E-02 | 0.583 | 3.6E-02 |
| LINC02733 | 1.272 | 4.1E-03 | 0.63 | 2.2E-02 | 0.549 | 4.4E-02 |
| LINC02774 | -1.156 | 6.1E-05 | -0.643 | 1.5E-03 | -0.61 | 2.7E-03 |
| LITAF | 0.909 | 3.0E-05 | 0.526 | 1.3E-03 | 0.352 | 3.4E-02 |
| LLGL2 | 1.048 | 2.2E-03 | 0.583 | 4.4E-03 | 0.458 | 2.6E-02 |
| LMF1-AS1 | 0.608 | 1.4E-03 | 0.298 | 9.5E-03 | 0.301 | 8.6E-03 |
| LNCAROD | 2.091 | 4.1E-05 | 1.216 | 2.9E-04 | 0.873 | 9.4E-03 |
| LNPK | -0.394 | 8.3E-05 | -0.261 | 1.5E-05 | -0.145 | 1.6E-02 |
| LRCH4 | 0.613 | 1.1E-03 | 0.32 | 3.1E-03 | 0.284 | 8.0E-03 |
| LRP2 | 1.767 | 7.3E-04 | 1.064 | 1.6E-03 | 0.703 | 4.1E-02 |
| LRP4 | 0.96 | 1.2E-03 | 0.517 | 1.0E-02 | 0.433 | 3.1E-02 |
| LRRC1 | 1.332 | 5.6E-05 | 0.681 | 4.2E-04 | 0.663 | 9.1E-04 |
| LRRC69 | 0.77 | 2.1E-03 | 0.383 | 1.1E-03 | 0.391 | 1.0E-03 |
| LRRK2-DT | -1.071 | 6.6E-03 | -0.501 | 4.2E-02 | -0.531 | 3.1E-02 |
| LRTM2 | -0.941 | 3.4E-04 | -0.409 | 4.4E-02 | -0.576 | 4.6E-03 |
| LUC7L2 | -0.555 | 2.7E-03 | -0.246 | 4.9E-02 | -0.332 | 7.3E-03 |
| LYRM9 | -1.046 | 1.9E-03 | -0.46 | 1.1E-02 | -0.658 | 2.9E-04 |
| LZTS2 | 0.549 | 2.5E-03 | 0.279 | 2.4E-02 | 0.262 | 3.4E-02 |
| MACROH2A2 | -0.391 | 2.3E-04 | -0.192 | 4.1E-03 | -0.214 | 3.5E-03 |
| MAD1L1 | 0.712 | 1.3E-05 | 0.393 | 1.9E-04 | 0.298 | 5.1E-03 |
| MAGED4/MAGEI | 2.825 | 3.4E-04 | 1.834 | 2.9E-03 | 1.605 | 1.7E-02 |
| MAGEH1 | -0.574 | 6.8E-03 | -0.312 | 1.9E-02 | -0.279 | 3.1E-02 |
| MAP1LC3A | -0.435 | 8.5E-03 | -0.245 | 1.2E-02 | -0.202 | 4.0E-02 |
| MAP2K2 | 0.299 | 1.0E-02 | 0.127 | 4.0E-02 | 0.164 | 9.3E-03 |
| MAP2K7 | 0.438 | 6.0E-03 | 0.225 | 2.4E-02 | 0.215 | 2.9E-02 |
| MAP4K3-DT | -0.552 | 1.2E-04 | -0.301 | 1.9E-02 | -0.267 | 3.8E-02 |
| MAP4K5 | 0.559 | 1.1E-02 | 0.297 | 1.1E-02 | 0.25 | 4.0E-02 |
| MAPK8 | -0.277 | 3.2E-03 | -0.136 | 3.2E-02 | -0.15 | 2.5E-02 |
| MAVS | 0.519 | 3.0E-03 | 0.324 | 1.1E-03 | 0.197 | 4.6E-02 |
| MBNL1 | 0.356 | 2.6E-03 | 0.176 | 4.8E-03 | 0.166 | 8.2E-03 |
| MCM3AP-AS1 | 0.333 | 1.0E-02 | 0.168 | 1.9E-02 | 0.156 | 2.8E-02 |
| MCM4 | -0.578 | 9.1E-03 | -0.307 | 6.1E-03 | -0.279 | 1.2E-02 |
| MCM7 | 0.771 | 6.8E-03 | 0.369 | 2.4E-02 | 0.414 | 1.0E-02 |
| MCPH1 | 0.381 | 2.0E-03 | 0.176 | 5.0E-02 | 0.196 | 3.0E-02 |
| MDH1B | -1.162 | 5.1E-06 | -0.596 | 3.1E-03 | -0.603 | 1.7E-03 |
| MEAF6 | -0.372 | 5.4E-03 | -0.168 | 3.7E-02 | -0.212 | 7.3E-03 |
| MED13L | 0.189 | 8.6E-03 | 0.081 | 1.7E-02 | 0.097 | 4.3E-03 |
| MED25 | 0.423 | 1.4E-03 | 0.183 | 4.3E-02 | 0.231 | 1.1E-02 |
| MEG9 | -1.309 | 8.0E-06 | -0.762 | 3.0E-04 | -0.551 | 9.3E-03 |
| METAP1D | -0.646 | 8.8E-04 | -0.299 | 2.3E-02 | -0.361 | 5.8E-03 |

| | | | | | | |
|---|---|---|---|---|---|---|
| MFN2 | -0.285 | 1.0E-03 | -0.133 | 2.2E-02 | -0.166 | 4.8E-03 |
| MFSD8 | -0.431 | 2.0E-03 | -0.24 | 2.0E-03 | -0.199 | 9.5E-03 |
| MICALL2 | 0.616 | 3.6E-03 | 0.308 | 2.7E-02 | 0.302 | 3.3E-02 |
| MID1IP1 | 0.974 | 7.7E-04 | 0.359 | 2.0E-02 | 0.597 | 1.1E-04 |
| MIPEP | -0.449 | 5.6E-03 | -0.232 | 2.8E-02 | -0.217 | 4.4E-02 |
| mir-1249 | 1.233 | 1.9E-03 | 0.577 | 2.0E-02 | 0.675 | 7.3E-03 |
| MIRLET7BHG | 0.55 | 3.3E-03 | 0.353 | 1.5E-04 | 0.189 | 3.9E-02 |
| MLKL | 1.036 | 1.1E-04 | 0.581 | 1.8E-03 | 0.398 | 3.4E-02 |
| MNAT1 | -0.38 | 1.8E-03 | -0.164 | 4.0E-02 | -0.22 | 5.3E-03 |
| MOB1A | 0.36 | 5.5E-04 | 0.179 | 2.2E-02 | 0.169 | 3.0E-02 |
| MOB3A | 0.478 | 3.2E-03 | 0.233 | 9.2E-03 | 0.241 | 4.9E-03 |
| MOG | 5.592 | 5.6E-06 | 2.88 | 5.2E-04 | 2.348 | 3.6E-03 |
| MPST | 1.003 | 4.8E-03 | 0.629 | 4.9E-04 | 0.376 | 3.6E-02 |
| MRPL15 | -0.348 | 9.8E-03 | -0.177 | 2.4E-02 | -0.18 | 2.1E-02 |
| MRPL33 | -0.354 | 9.2E-03 | -0.224 | 2.2E-03 | -0.144 | 4.8E-02 |
| MRPS30-DT | -0.583 | 5.7E-04 | -0.281 | 3.0E-02 | -0.318 | 1.4E-02 |
| MS4A14 | 1.406 | 2.0E-05 | 0.747 | 7.6E-03 | 0.644 | 2.2E-02 |
| MS4A6E | 1.201 | 5.4E-04 | 0.558 | 3.3E-02 | 0.642 | 1.4E-02 |
| MS4A7 | 1.254 | 2.5E-04 | 0.615 | 1.9E-02 | 0.641 | 1.5E-02 |
| MS4A8 | -1.658 | 3.5E-03 | -0.918 | 6.0E-03 | -0.847 | 1.1E-02 |
| MT1E | 1.193 | 5.8E-06 | 0.634 | 5.0E-04 | 0.552 | 2.7E-03 |
| MT1F | 1.57 | 1.5E-09 | 0.982 | 9.7E-06 | 0.593 | 7.6E-03 |
| MT1G | 1.476 | 1.7E-05 | 0.798 | 1.3E-03 | 0.642 | 1.2E-02 |
| MT1JP | 4.59 | 5.7E-04 | 1.934 | 6.2E-03 | 2.037 | 4.7E-03 |
| MT1L | 1.799 | 4.4E-04 | 0.786 | 1.6E-02 | 0.896 | 5.7E-03 |
| MTMR9 | -0.438 | 4.8E-07 | -0.238 | 2.2E-04 | -0.211 | 3.5E-04 |
| MTX3 | -0.358 | 2.5E-04 | -0.158 | 1.8E-02 | -0.217 | 7.5E-04 |
| MXRA8 | -0.725 | 1.4E-03 | -0.353 | 1.0E-02 | -0.388 | 6.0E-03 |
| MYLK-AS1 | 0.551 | 8.4E-03 | 0.271 | 3.4E-02 | 0.283 | 2.8E-02 |
| MYO9B | 0.652 | 3.6E-04 | 0.335 | 8.5E-03 | 0.312 | 1.4E-02 |
| NACC2 | 0.863 | 8.0E-05 | 0.572 | 7.7E-07 | 0.3 | 8.8E-03 |
| NAP1L5 | -0.914 | 5.7E-05 | -0.565 | 8.5E-04 | -0.354 | 3.8E-02 |
| NBEAL2 | 0.416 | 1.8E-02 | 0.204 | 4.9E-02 | 0.202 | 4.5E-02 |
| NBPF10 (include: | -0.378 | 1.1E-02 | -0.321 | 9.7E-04 | 0.307 | 3.2E-02 |
| NCLP1 | 1.001 | 1.4E-03 | 0.543 | 8.5E-03 | 0.426 | 4.0E-02 |
| NCMAP | 2.5 | 5.5E-06 | 1.092 | 2.4E-03 | 1.352 | 1.4E-04 |
| NCOR2 | 0.387 | 1.9E-03 | 0.21 | 1.2E-02 | 0.17 | 3.5E-02 |
| NCR3LG1 | -0.602 | 3.2E-04 | -0.351 | 4.9E-03 | -0.26 | 3.8E-02 |
| NDFIP1 | -0.482 | 1.5E-03 | -0.249 | 9.1E-03 | -0.235 | 1.3E-02 |
| NDRG1 | 1.011 | 2.8E-04 | 0.559 | 1.7E-03 | 0.449 | 1.2E-02 |
| NDUFA5 | -0.568 | 7.6E-05 | -0.309 | 8.2E-04 | -0.263 | 6.0E-03 |
| NDUFAF6 | 0.518 | 3.1E-03 | 0.272 | 2.9E-03 | 0.226 | 1.3E-02 |
| NDUFB5 | -0.525 | 1.5E-03 | -0.261 | 8.0E-03 | -0.275 | 4.1E-03 |
| NEAT1 | 1.478 | 1.8E-07 | 0.599 | 4.1E-03 | 0.86 | 3.9E-05 |
| NFASC | 0.525 | 5.8E-03 | 0.264 | 1.2E-02 | 0.253 | 1.9E-02 |
| NFATC1 | 1.44 | 3.9E-06 | 0.731 | 4.1E-04 | 0.662 | 1.2E-03 |
| NFATC2 | 1.248 | 4.9E-12 | 0.444 | 6.3E-03 | 0.772 | 1.8E-06 |
| NGFR | 2.156 | 1.8E-03 | 1.145 | 2.1E-02 | 1.116 | 2.4E-02 |
| NKX6-2 | 1.75 | 1.2E-05 | 1.08 | 7.0E-07 | 0.672 | 2.0E-03 |
| NMD3 | -0.546 | 8.8E-04 | -0.283 | 8.3E-03 | -0.277 | 9.1E-03 |

| | | | | | | |
|---|---|---|---|---|---|---|
| NMUR2 | 1.751 | 3.1E-03 | 0.9 | 2.2E-02 | 0.872 | 3.0E-02 |
| NORAD | -0.367 | 5.7E-05 | -0.185 | 1.8E-03 | -0.192 | 8.9E-04 |
| NPRL3 | 0.349 | 1.8E-02 | 0.178 | 1.3E-02 | 0.157 | 2.8E-02 |
| NPY6R | 0.86 | 2.2E-03 | 0.446 | 9.9E-03 | 0.386 | 3.1E-02 |
| NRIP2 | 0.554 | 5.0E-03 | 0.312 | 5.6E-03 | 0.232 | 4.6E-02 |
| NRSN2 | -0.591 | 1.3E-03 | -0.246 | 3.9E-02 | -0.369 | 1.9E-03 |
| NSG1 | -0.7 | 1.3E-04 | -0.377 | 7.9E-03 | -0.335 | 1.8E-02 |
| NUDT21 | -0.429 | 9.1E-04 | -0.267 | 9.6E-04 | -0.167 | 4.1E-02 |
| NUP42 | -0.482 | 1.7E-03 | -0.24 | 2.8E-02 | -0.257 | 1.8E-02 |
| OLMALINC | 1.365 | 6.9E-08 | 0.779 | 4.2E-06 | 0.585 | 9.8E-04 |
| OPTC | 1.507 | 2.7E-03 | 0.863 | 2.0E-02 | 0.852 | 1.9E-02 |
| OR10G4 | -4.435 | 7.5E-04 | -1.725 | 2.2E-02 | -2.088 | 8.0E-03 |
| OR2AG2 | -1.027 | 1.8E-03 | -0.575 | 2.7E-02 | -0.536 | 3.8E-02 |
| OR7A5 | 3.438 | 2.4E-04 | 1.554 | 4.8E-03 | 1.916 | 7.8E-04 |
| OR7C1 | 2.721 | 1.3E-03 | 1.092 | 9.9E-03 | 1.425 | 9.0E-04 |
| OR7E161P | -1.515 | 2.5E-03 | -0.757 | 2.7E-02 | -0.757 | 2.5E-02 |
| ORAI2 | 0.77 | 1.8E-05 | 0.372 | 2.7E-03 | 0.381 | 2.1E-03 |
| OTUD7B | 0.789 | 5.5E-04 | 0.388 | 1.3E-02 | 0.393 | 1.2E-02 |
| P2RX7 | 0.678 | 1.2E-03 | 0.385 | 5.5E-04 | 0.283 | 1.1E-02 |
| PACRGL | -0.277 | 4.5E-03 | -0.158 | 3.3E-03 | -0.134 | 7.4E-03 |
| PACS2 | 0.638 | 7.5E-06 | 0.372 | 9.3E-05 | 0.26 | 6.5E-03 |
| PAK1 | -0.817 | 2.7E-05 | -0.465 | 3.8E-03 | -0.369 | 2.0E-02 |
| PAK2 | 0.437 | 4.9E-04 | 0.176 | 1.9E-02 | 0.248 | 8.3E-04 |
| PAK6 | -0.773 | 3.0E-04 | -0.406 | 2.8E-02 | -0.394 | 3.8E-02 |
| PANK2-AS1 | 1.282 | 2.2E-03 | 0.603 | 1.8E-02 | 0.664 | 1.0E-02 |
| PANK3 | -0.268 | 6.9E-05 | -0.159 | 1.6E-04 | -0.12 | 4.2E-03 |
| PARVG | 1.163 | 1.2E-06 | 0.423 | 1.3E-02 | 0.718 | 5.6E-05 |
| PAX6 | 0.556 | 2.6E-06 | 0.264 | 1.8E-02 | 0.282 | 1.4E-02 |
| PAXX | 0.601 | 1.7E-02 | 0.335 | 1.1E-02 | 0.26 | 4.5E-02 |
| PCA3 | 1.13 | 1.3E-03 | 0.642 | 3.2E-03 | 0.486 | 3.0E-02 |
| PCDH1 | -0.429 | 7.9E-04 | -0.204 | 4.0E-02 | -0.237 | 1.7E-02 |
| PCYOX1L | -1.009 | 3.4E-05 | -0.539 | 2.0E-03 | -0.523 | 2.6E-03 |
| PDE10A | -0.899 | 3.4E-08 | -0.399 | 3.8E-03 | -0.508 | 1.4E-04 |
| PFKFB4 | 0.527 | 2.0E-03 | 0.292 | 1.3E-03 | 0.22 | 1.4E-02 |
| PFN2 | -0.623 | 1.5E-05 | -0.332 | 8.7E-03 | -0.297 | 1.6E-02 |
| PGAP4 | -0.578 | 9.0E-05 | -0.307 | 2.6E-02 | -0.282 | 3.3E-02 |
| PGLS-DT | 0.664 | 2.3E-04 | 0.446 | 1.5E-07 | 0.212 | 1.2E-02 |
| PGPEP1L | -0.94 | 4.8E-04 | -0.511 | 5.3E-03 | -0.426 | 2.0E-02 |
| PGRMC1 | -0.448 | 5.6E-04 | -0.266 | 1.2E-03 | -0.194 | 1.9E-02 |
| PGS1 | 0.459 | 7.1E-07 | 0.27 | 6.9E-05 | 0.177 | 8.7E-03 |
| PHACTR3 | 0.434 | 6.8E-03 | 0.189 | 3.5E-02 | 0.237 | 7.4E-03 |
| PHF19 | 1.063 | 1.0E-05 | 0.54 | 1.3E-03 | 0.503 | 2.8E-03 |
| PHF21A | 0.35 | 2.3E-04 | 0.123 | 4.7E-02 | 0.216 | 3.9E-04 |
| PIAS4 | 0.69 | 4.5E-06 | 0.352 | 6.3E-06 | 0.323 | 3.6E-05 |
| PICALM | 0.549 | 6.7E-04 | 0.307 | 3.2E-04 | 0.23 | 1.0E-02 |
| PIEZO1 | 0.849 | 4.5E-04 | 0.386 | 5.1E-03 | 0.479 | 3.3E-04 |
| PIK3AP1 | 1.243 | 6.8E-05 | 0.605 | 4.4E-02 | 0.606 | 4.2E-02 |
| PIK3CD | 0.439 | 1.2E-02 | 0.244 | 1.8E-02 | 0.197 | 4.7E-02 |
| PIPOX | -0.639 | 7.2E-07 | -0.294 | 5.6E-03 | -0.36 | 5.2E-04 |
| PJA2 | -0.476 | 2.0E-03 | -0.265 | 1.4E-02 | -0.211 | 4.6E-02 |

| | | | | | | |
|---|---|---|---|---|---|---|
| PKN2-AS1 | -0.717 | 5.3E-05 | -0.408 | 4.3E-03 | -0.322 | 2.4E-02 |
| PLEKHA3 | -0.331 | 3.2E-03 | -0.191 | 6.9E-03 | -0.151 | 2.7E-02 |
| PLEKHA7 | 0.762 | 2.3E-05 | 0.358 | 3.1E-02 | 0.401 | 1.7E-02 |
| PLEKHB1 | 0.716 | 1.1E-03 | 0.413 | 1.3E-03 | 0.31 | 1.9E-02 |
| PLEKHB2 | -0.475 | 4.2E-04 | -0.283 | 9.4E-04 | -0.203 | 1.7E-02 |
| PLIN3 | 1.365 | 1.1E-06 | 0.605 | 7.8E-03 | 0.735 | 1.2E-03 |
| PLOD3 | 1.047 | 6.6E-04 | 0.669 | 7.7E-06 | 0.332 | 2.7E-02 |
| PLPP4 | 0.99 | 3.6E-04 | 0.45 | 8.4E-03 | 0.534 | 9.1E-04 |
| PLXNB1 | 0.959 | 1.4E-05 | 0.544 | 4.3E-05 | 0.414 | 1.8E-03 |
| PMS2P4 | -0.63 | 1.3E-03 | -0.275 | 3.7E-02 | -0.375 | 4.4E-03 |
| PNMA3 | -0.837 | 9.2E-03 | -0.433 | 4.8E-02 | -0.49 | 2.5E-02 |
| PNMA8C | -0.929 | 1.0E-04 | -0.476 | 9.3E-03 | -0.454 | 1.1E-02 |
| POGK | 0.37 | 2.5E-02 | 0.178 | 2.9E-02 | 0.183 | 2.5E-02 |
| POLD1 | 1.114 | 3.0E-07 | 0.528 | 8.9E-05 | 0.558 | 2.2E-05 |
| POLR2K | -0.602 | 9.3E-05 | -0.371 | 1.4E-04 | -0.243 | 1.1E-02 |
| PPEF1 | -1.144 | 4.3E-04 | -0.572 | 1.0E-02 | -0.591 | 6.9E-03 |
| PPFIBP2 | 1.019 | 2.4E-03 | 0.559 | 2.0E-03 | 0.46 | 1.3E-02 |
| PPIP5K2 | -0.32 | 6.4E-03 | -0.193 | 6.7E-04 | -0.137 | 2.0E-02 |
| PPP1R2 | -0.563 | 8.6E-04 | -0.354 | 3.6E-05 | -0.225 | 8.8E-03 |
| PPP1R3B | 0.753 | 1.1E-03 | 0.356 | 4.5E-02 | 0.372 | 4.6E-02 |
| PPP4R2 | 0.467 | 5.8E-03 | 0.227 | 8.9E-03 | 0.222 | 1.1E-02 |
| PPT1 | -0.379 | 1.8E-03 | -0.169 | 2.5E-02 | -0.218 | 1.1E-03 |
| PREX1 | 0.912 | 1.2E-05 | 0.547 | 3.3E-04 | 0.354 | 2.1E-02 |
| PRKAR1A | -0.353 | 1.4E-03 | -0.222 | 1.9E-03 | -0.141 | 4.2E-02 |
| PRKX | 1.466 | 5.6E-09 | 0.752 | 1.1E-04 | 0.681 | 5.9E-04 |
| PROC | 1.076 | 9.8E-03 | 0.522 | 4.0E-02 | 0.513 | 4.4E-02 |
| PRRG1 | 0.751 | 3.5E-04 | 0.377 | 9.2E-04 | 0.362 | 2.6E-03 |
| PSG3 | -1.317 | 2.1E-03 | -0.562 | 2.3E-02 | -0.817 | 7.5E-04 |
| PSMA3 | 0.365 | 1.9E-03 | 0.155 | 9.2E-03 | 0.2 | 9.5E-04 |
| PSMG4 | 0.77 | 3.1E-04 | 0.39 | 1.6E-02 | 0.366 | 2.2E-02 |
| PTP4A2 | 0.596 | 7.2E-03 | 0.303 | 1.8E-03 | 0.263 | 8.9E-03 |
| PTPRC | 1.197 | 1.0E-05 | 0.551 | 2.9E-03 | 0.585 | 1.3E-03 |
| PUM3 | -0.559 | 1.5E-05 | -0.32 | 1.8E-04 | -0.252 | 2.6E-03 |
| QDPR | 1.217 | 3.6E-04 | 0.683 | 1.5E-04 | 0.536 | 3.4E-03 |
| R3HCC1 | 0.706 | 1.6E-03 | 0.343 | 2.8E-02 | 0.361 | 2.1E-02 |
| RALGAPA2 | 0.452 | 6.0E-05 | 0.215 | 2.2E-03 | 0.225 | 1.4E-03 |
| RALGDS | 0.719 | 1.0E-03 | 0.371 | 2.5E-02 | 0.34 | 3.7E-02 |
| RASA3 | 0.377 | 1.4E-03 | 0.168 | 2.7E-02 | 0.197 | 9.6E-03 |
| RASSF2 | 0.914 | 4.4E-03 | 0.532 | 4.8E-03 | 0.384 | 4.9E-02 |
| RAVER1 | 0.355 | 1.4E-02 | 0.18 | 3.4E-02 | 0.173 | 4.2E-02 |
| RBFOX2 | -0.347 | 5.9E-04 | -0.203 | 5.3E-03 | -0.155 | 3.4E-02 |
| RBM19 | 0.338 | 2.2E-02 | 0.163 | 4.6E-02 | 0.168 | 3.9E-02 |
| RELA | 0.845 | 6.7E-05 | 0.381 | 5.1E-03 | 0.448 | 9.3E-04 |
| RENBP | 1.241 | 1.9E-05 | 0.751 | 2.1E-06 | 0.502 | 1.8E-03 |
| RESF1 | 0.427 | 3.0E-03 | 0.199 | 2.6E-02 | 0.214 | 1.5E-02 |
| RFTN2 | 1.02 | 1.9E-03 | 0.617 | 5.0E-04 | 0.396 | 2.7E-02 |
| RGCC | 1.168 | 4.8E-05 | 0.496 | 1.3E-02 | 0.664 | 7.5E-04 |
| RGS17 | -0.624 | 2.9E-04 | -0.231 | 2.8E-02 | -0.411 | 8.9E-05 |
| RGS18 | 1.463 | 1.1E-03 | 0.746 | 7.4E-03 | 0.674 | 1.5E-02 |
| RHBDF2 | 1.467 | 2.9E-08 | 0.771 | 5.8E-04 | 0.625 | 4.2E-03 |

| | | | | | | |
|---|---|---|---|---|---|---|
| RHOXF1P3 | -0.833 | 2.1E-04 | -0.438 | 3.3E-03 | -0.383 | 9.4E-03 |
| RIC3 | -0.289 | 3.5E-03 | -0.16 | 9.5E-03 | -0.139 | 2.6E-02 |
| RIN2 | 0.74 | 4.2E-06 | 0.45 | 7.9E-06 | 0.274 | 1.2E-02 |
| RNF121 | -0.283 | 9.1E-04 | -0.165 | 3.4E-03 | -0.124 | 2.8E-02 |
| RNF144A | 0.629 | 5.3E-04 | 0.34 | 4.4E-03 | 0.284 | 1.9E-02 |
| RNF166 | 0.898 | 4.5E-08 | 0.577 | 4.4E-07 | 0.305 | 6.7E-03 |
| RNF187 | -0.482 | 1.8E-03 | -0.256 | 2.6E-02 | -0.237 | 4.0E-02 |
| RNF216 | 0.305 | 6.5E-05 | 0.118 | 1.7E-02 | 0.173 | 4.5E-04 |
| RNF216-IT1 | 0.536 | 1.4E-04 | 0.184 | 4.6E-02 | 0.352 | 1.2E-04 |
| RNF34 | -0.26 | 6.2E-04 | -0.146 | 1.0E-03 | -0.123 | 8.7E-03 |
| RNMT | -0.434 | 2.7E-04 | -0.244 | 5.4E-05 | -0.206 | 8.7E-04 |
| RPGRIP1L | -0.581 | 1.5E-05 | -0.247 | 3.7E-02 | -0.349 | 2.4E-03 |
| RPL15 | -0.36 | 1.7E-03 | -0.195 | 7.4E-03 | -0.174 | 1.8E-02 |
| RPS10P7 | 0.602 | 3.8E-04 | 0.289 | 3.5E-02 | 0.293 | 3.4E-02 |
| RPS6KA1 | 1.107 | 4.6E-06 | 0.489 | 7.9E-03 | 0.553 | 2.5E-03 |
| RUNX1 | 1.044 | 1.2E-09 | 0.565 | 1.3E-05 | 0.474 | 2.6E-04 |
| RXRA | 0.685 | 2.9E-04 | 0.331 | 1.7E-02 | 0.343 | 1.3E-02 |
| SAFB | 0.367 | 5.2E-05 | 0.149 | 3.8E-03 | 0.206 | 6.2E-05 |
| SASH1 | 0.607 | 1.6E-04 | 0.348 | 1.9E-03 | 0.246 | 2.7E-02 |
| SCAMP2 | 0.645 | 3.8E-05 | 0.344 | 1.3E-03 | 0.287 | 7.9E-03 |
| SCMH1 | -0.279 | 1.7E-03 | -0.138 | 3.0E-02 | -0.152 | 1.6E-02 |
| SCOC | -0.55 | 2.8E-06 | -0.286 | 6.2E-04 | -0.275 | 5.7E-04 |
| SDCCAG8 | 1.019 | 7.5E-10 | 0.556 | 2.7E-02 | 0.416 | 4.9E-02 |
| SEC22A | -0.283 | 5.8E-04 | -0.13 | 1.5E-02 | -0.163 | 4.3E-03 |
| SEPTIN8 | 0.654 | 6.9E-04 | 0.355 | 8.7E-04 | 0.298 | 5.1E-03 |
| SERINC1 | -0.342 | 1.1E-02 | -0.177 | 1.8E-02 | -0.175 | 2.0E-02 |
| SEZ6 | -0.768 | 9.0E-08 | -0.373 | 1.4E-02 | -0.41 | 6.3E-03 |
| SF3B1 | 0.197 | 2.6E-02 | 0.093 | 4.7E-02 | 0.094 | 4.8E-02 |
| SGCA | 1.328 | 1.5E-03 | 0.622 | 1.9E-02 | 0.728 | 5.0E-03 |
| SH2D5 | -0.822 | 1.1E-03 | -0.424 | 4.5E-02 | -0.42 | 4.8E-02 |
| SH2D6 | 2.026 | 2.7E-08 | 0.901 | 4.6E-04 | 1.13 | 1.7E-05 |
| SHKBP1 | 0.845 | 2.3E-03 | 0.464 | 3.7E-03 | 0.376 | 1.8E-02 |
| SHMT1 | 0.919 | 6.4E-06 | 0.41 | 4.7E-03 | 0.498 | 6.2E-04 |
| SIPA1 | 1.067 | 1.1E-08 | 0.608 | 9.1E-05 | 0.426 | 5.2E-03 |
| SIRPB2 | 1.721 | 1.9E-05 | 0.718 | 5.7E-03 | 0.838 | 1.2E-03 |
| SLA | 1.278 | 6.0E-06 | 0.542 | 1.3E-02 | 0.653 | 2.4E-03 |
| SLC16A14 | -0.778 | 2.9E-04 | -0.328 | 1.7E-02 | -0.46 | 8.7E-04 |
| SLC16A6 | -1.384 | 4.7E-04 | -0.691 | 2.2E-02 | -0.724 | 1.8E-02 |
| SLC22A23 | 0.828 | 1.4E-06 | 0.467 | 2.1E-06 | 0.353 | 5.4E-04 |
| SLC2A5 | 1.691 | 1.8E-07 | 0.511 | 4.8E-02 | 1.106 | 1.3E-05 |
| SLC35B4 | -0.409 | 3.3E-04 | -0.218 | 6.3E-03 | -0.208 | 4.2E-03 |
| SLC38A10 | 0.469 | 1.2E-05 | 0.237 | 1.8E-03 | 0.226 | 2.6E-03 |
| SLC39A11 | 1.126 | 2.1E-04 | 0.51 | 2.5E-03 | 0.616 | 2.8E-04 |
| SLC5A11 | 1.784 | 6.5E-04 | 0.89 | 1.6E-02 | 0.923 | 1.4E-02 |
| SLC6A9 | 1.364 | 1.8E-06 | 0.691 | 3.0E-04 | 0.677 | 5.0E-04 |
| SLC9A9 | 0.582 | 6.7E-04 | 0.31 | 1.0E-02 | 0.256 | 3.5E-02 |
| SMC4 | 1.172 | 1.1E-03 | 0.73 | 1.7E-04 | 0.441 | 2.9E-02 |
| SMG5 | 0.341 | 6.0E-03 | 0.16 | 2.1E-02 | 0.174 | 1.3E-02 |
| SMIM10L2B | -0.894 | 3.4E-04 | -0.473 | 3.2E-02 | -0.49 | 2.6E-02 |
| SMPD4 | 0.44 | 3.4E-05 | 0.215 | 2.6E-04 | 0.218 | 1.7E-04 |

| | | | | | | |
|---|---|---|---|---|---|---|
| SNORD17 | 0.584 | 3.5E-03 | 0.311 | 9.6E-03 | 0.247 | 3.9E-02 |
| SNRK | -0.304 | 3.4E-04 | -0.155 | 8.1E-04 | -0.157 | 5.1E-04 |
| SNU13 | -0.466 | 2.0E-03 | -0.268 | 9.4E-03 | -0.213 | 3.5E-02 |
| SNX20 | 1.587 | 1.2E-03 | 0.696 | 3.5E-02 | 0.794 | 1.3E-02 |
| SOWAHB | -1.084 | 9.2E-03 | -0.588 | 1.7E-02 | -0.578 | 1.9E-02 |
| SPATA13 | 0.952 | 7.8E-04 | 0.44 | 3.1E-02 | 0.485 | 2.1E-02 |
| SPCS1 | -0.373 | 6.7E-04 | -0.158 | 4.2E-02 | -0.228 | 3.0E-03 |
| SPIN2A/SPIN2B | -0.823 | 2.0E-02 | -0.619 | 4.5E-03 | -0.395 | 3.7E-02 |
| SPN | 1.388 | 7.8E-03 | 0.652 | 3.4E-02 | 0.617 | 4.6E-02 |
| SPSB1 | 1.123 | 1.9E-06 | 0.622 | 1.8E-03 | 0.463 | 2.1E-02 |
| SPTSSB | -0.91 | 8.4E-04 | -0.485 | 1.9E-02 | -0.444 | 3.2E-02 |
| SRGAP1 | 0.536 | 7.4E-06 | 0.279 | 7.4E-04 | 0.246 | 2.8E-03 |
| ST6GALNAC3 | 0.632 | 1.7E-05 | 0.299 | 2.0E-02 | 0.32 | 1.5E-02 |
| ST7L | 0.291 | 8.6E-04 | 0.132 | 1.6E-02 | 0.148 | 8.2E-03 |
| STAG2 | 0.288 | 6.2E-03 | 0.137 | 2.7E-02 | 0.142 | 2.9E-02 |
| STARD3 | 0.623 | 1.1E-04 | 0.278 | 1.6E-03 | 0.336 | 1.3E-04 |
| STON1-GTF2A1I | 1.185 | 1.3E-06 | 0.418 | 2.4E-02 | 0.771 | 4.1E-05 |
| STPG4 | 0.566 | 5.0E-03 | 0.288 | 2.7E-02 | 0.267 | 4.0E-02 |
| STS | -0.669 | 9.0E-03 | -0.285 | 1.2E-02 | -0.415 | 2.1E-04 |
| STUM | -0.937 | 1.5E-09 | -0.464 | 4.2E-03 | -0.493 | 2.4E-03 |
| SUB1 | -0.755 | 2.8E-06 | -0.447 | 1.9E-03 | -0.306 | 3.4E-02 |
| SUN2 | 0.961 | 4.4E-04 | 0.535 | 7.2E-04 | 0.421 | 8.4E-03 |
| SUN3 | -1.648 | 3.0E-03 | -0.604 | 4.7E-02 | -0.979 | 1.5E-03 |
| SUSD5 | -0.441 | 3.5E-03 | -0.222 | 2.9E-02 | -0.237 | 2.7E-02 |
| SWAP70 | 0.368 | 6.3E-04 | 0.165 | 3.2E-02 | 0.189 | 1.2E-02 |
| SYK | 1.494 | 1.3E-05 | 0.759 | 1.4E-03 | 0.655 | 5.6E-03 |
| SYT11 | -0.352 | 1.6E-03 | -0.188 | 4.1E-03 | -0.18 | 6.3E-03 |
| SYTL2 | -0.467 | 1.2E-05 | -0.252 | 1.5E-02 | -0.225 | 3.2E-02 |
| SYTL4 | 1.389 | 6.8E-04 | 0.666 | 1.5E-02 | 0.721 | 8.2E-03 |
| TANC1 | 0.853 | 1.6E-05 | 0.502 | 4.3E-05 | 0.332 | 1.5E-02 |
| TARBP1 | -0.658 | 9.5E-04 | -0.276 | 4.5E-03 | -0.404 | 4.1E-05 |
| TASL | 1.992 | 1.7E-03 | 0.885 | 1.8E-02 | 0.949 | 1.0E-02 |
| TASOR | 0.261 | 1.9E-02 | 0.117 | 4.0E-02 | 0.133 | 2.4E-02 |
| TASP1 | -0.622 | 2.5E-06 | -0.334 | 7.4E-04 | -0.3 | 2.3E-03 |
| TBC1D9 | -0.653 | 2.8E-04 | -0.39 | 8.6E-05 | -0.273 | 5.6E-03 |
| TBL1X | 0.526 | 4.3E-04 | 0.272 | 2.5E-02 | 0.241 | 4.6E-02 |
| TBX6 | 1.618 | 7.8E-05 | 0.771 | 5.3E-03 | 0.796 | 4.1E-03 |
| TBXAS1 | 1.3 | 3.2E-06 | 0.778 | 4.2E-05 | 0.498 | 7.3E-03 |
| TCEAL7 | -0.461 | 4.5E-03 | -0.233 | 1.4E-02 | -0.244 | 1.0E-02 |
| TCIRG1 | 0.927 | 6.5E-05 | 0.537 | 5.0E-03 | 0.383 | 4.7E-02 |
| TCTE1 | -0.684 | 7.8E-04 | -0.345 | 2.3E-02 | -0.373 | 1.4E-02 |
| TDG | -0.374 | 2.5E-02 | -0.193 | 3.1E-02 | -0.192 | 3.1E-02 |
| TECTA | -0.679 | 4.2E-04 | -0.439 | 2.0E-05 | -0.259 | 1.2E-02 |
| TEPSIN | 0.532 | 1.4E-02 | 0.26 | 2.3E-02 | 0.275 | 1.7E-02 |
| TEX22 | 0.457 | 8.5E-03 | 0.213 | 1.3E-02 | 0.233 | 8.6E-03 |
| TFEB | 1.141 | 1.7E-04 | 0.635 | 8.0E-04 | 0.499 | 8.9E-03 |
| TFEC | 1.087 | 7.5E-05 | 0.499 | 1.9E-02 | 0.538 | 1.0E-02 |
| THBS2 | 0.924 | 1.2E-03 | 0.386 | 2.3E-02 | 0.561 | 1.2E-03 |
| THCAT155 | -1.155 | 2.3E-05 | -0.633 | 1.2E-02 | -0.558 | 2.6E-02 |
| THEMIS2 | 1.005 | 2.9E-05 | 0.454 | 1.6E-02 | 0.515 | 6.0E-03 |

| | | | | | | |
|---|---|---|---|---|---|---|
| TIMM44 | 0.685 | 3.6E-05 | 0.233 | 4.3E-02 | 0.429 | 1.0E-04 |
| TLDC2 | 0.828 | 4.6E-05 | 0.378 | 9.3E-03 | 0.434 | 2.9E-03 |
| TLE4 | 0.751 | 5.2E-04 | 0.432 | 3.7E-04 | 0.305 | 1.5E-02 |
| TLN1 | 0.437 | 4.6E-04 | 0.222 | 2.8E-02 | 0.211 | 3.9E-02 |
| TLR1 | 1.339 | 1.1E-06 | 0.611 | 2.4E-03 | 0.659 | 7.6E-04 |
| TLR5 | 1.183 | 2.1E-05 | 0.438 | 7.7E-03 | 0.722 | 8.9E-06 |
| TLR7 | 1.537 | 2.2E-03 | 0.825 | 1.9E-02 | 0.765 | 3.0E-02 |
| TLR8 | 1.8 | 1.4E-04 | 0.76 | 2.7E-02 | 0.91 | 2.7E-03 |
| TMC6 | 1.272 | 2.6E-04 | 0.736 | 7.8E-04 | 0.523 | 1.8E-02 |
| TMC8 | 1.557 | 5.7E-05 | 0.787 | 4.8E-03 | 0.651 | 1.9E-02 |
| TMCC2 | 0.786 | 2.0E-04 | 0.423 | 2.6E-03 | 0.357 | 1.2E-02 |
| TMEM132E | -0.518 | 1.0E-02 | -0.271 | 4.4E-02 | -0.269 | 4.5E-02 |
| TMEM14A | -0.58 | 5.8E-03 | -0.252 | 3.4E-02 | -0.345 | 3.3E-03 |
| TMEM53 | 0.597 | 3.8E-03 | 0.365 | 1.2E-03 | 0.228 | 4.5E-02 |
| TOB2 | 0.717 | 1.0E-03 | 0.383 | 1.9E-03 | 0.322 | 9.1E-03 |
| TP53I3 | 0.69 | 2.8E-04 | 0.427 | 3.0E-04 | 0.245 | 4.0E-02 |
| TPBGL | -1.515 | 1.7E-03 | -0.897 | 2.1E-03 | -0.675 | 2.2E-02 |
| TPRN | 0.808 | 2.0E-03 | 0.495 | 1.1E-03 | 0.314 | 3.7E-02 |
| TRDN | 1.284 | 5.1E-04 | 0.62 | 3.8E-02 | 0.676 | 2.0E-02 |
| TRIM3 | -0.346 | 1.6E-03 | -0.205 | 3.8E-03 | -0.159 | 2.7E-02 |
| TRIM33 | -0.22 | 7.5E-03 | -0.105 | 1.3E-02 | -0.124 | 6.1E-03 |
| TRIM36 | -0.881 | 9.0E-05 | -0.433 | 9.4E-03 | -0.454 | 5.9E-03 |
| TRIM62 | 0.563 | 5.4E-03 | 0.292 | 9.3E-03 | 0.261 | 1.9E-02 |
| TRIM8 | 0.498 | 2.2E-03 | 0.3 | 1.5E-03 | 0.191 | 4.5E-02 |
| TRNAU1AP | 0.416 | 7.1E-03 | 0.186 | 3.2E-02 | 0.218 | 1.1E-02 |
| TRRAP | 0.279 | 2.9E-04 | 0.128 | 7.5E-03 | 0.14 | 2.5E-03 |
| TSC22D3 | 0.781 | 1.8E-04 | 0.411 | 2.4E-03 | 0.362 | 6.8E-03 |
| TSEN15 | 0.649 | 6.1E-05 | 0.282 | 3.7E-03 | 0.356 | 3.7E-04 |
| TSFM | -0.445 | 4.6E-03 | -0.233 | 8.0E-03 | -0.227 | 1.1E-02 |
| TSPO | 0.948 | 6.7E-05 | 0.419 | 3.5E-02 | 0.499 | 1.2E-02 |
| TSPYL5 | -0.612 | 1.7E-03 | -0.33 | 1.2E-02 | -0.287 | 3.0E-02 |
| TTC5 | -0.787 | 1.8E-04 | -0.395 | 1.6E-03 | -0.399 | 9.8E-04 |
| TTYH2 | 1.22 | 2.4E-03 | 0.694 | 7.2E-03 | 0.519 | 4.6E-02 |
| TUBA1B | -0.623 | 9.5E-06 | -0.258 | 3.7E-02 | -0.37 | 2.3E-03 |
| UBE2R2 | 0.353 | 6.0E-05 | 0.148 | 5.3E-04 | 0.194 | 2.6E-05 |
| UBFD1 | -0.377 | 1.2E-03 | -0.224 | 3.6E-03 | -0.164 | 3.5E-02 |
| UBLCP1 | -0.458 | 1.4E-04 | -0.259 | 3.8E-04 | -0.209 | 9.9E-03 |
| UBXN2A | 0.466 | 1.7E-03 | 0.235 | 5.1E-03 | 0.215 | 9.9E-03 |
| UCKL1 | 0.786 | 2.7E-07 | 0.461 | 6.2E-07 | 0.318 | 5.3E-04 |
| UHRF1 | 1.234 | 1.3E-04 | 0.541 | 5.6E-03 | 0.675 | 4.4E-04 |
| UNKL | 0.558 | 5.7E-04 | 0.282 | 4.3E-03 | 0.265 | 4.5E-03 |
| USP11 | -0.614 | 2.4E-03 | -0.295 | 2.6E-02 | -0.328 | 1.3E-02 |
| USP47 | 0.407 | 1.2E-04 | 0.161 | 3.6E-03 | 0.232 | 6.8E-05 |
| USP5 | -0.415 | 4.2E-03 | -0.22 | 3.2E-02 | -0.212 | 3.9E-02 |
| UTP25 | -0.342 | 5.2E-04 | -0.204 | 1.3E-03 | -0.147 | 1.9E-02 |
| VAC14 | 0.284 | 1.5E-02 | 0.134 | 2.4E-02 | 0.145 | 1.3E-02 |
| VAPB | -0.329 | 4.3E-03 | -0.223 | 2.7E-04 | -0.121 | 4.6E-02 |
| VAV1 | 1.298 | 8.8E-05 | 0.663 | 9.5E-03 | 0.557 | 2.8E-02 |
| VEZF1 | 0.753 | 3.8E-03 | 0.392 | 2.2E-03 | 0.352 | 6.5E-03 |
| VEZT | -0.339 | 3.3E-04 | -0.186 | 2.1E-04 | -0.163 | 1.3E-03 |

| | | | | | | |
|---|---|---|---|---|---|---|
| VSIR | 1.037 | 2.8E-05 | 0.501 | 1.4E-02 | 0.522 | 8.2E-03 |
| WDR64 | 1.632 | 1.9E-03 | 0.78 | 4.7E-02 | 1.092 | 4.2E-03 |
| WDR86-AS1 | -2.324 | 5.7E-03 | -1.22 | 1.7E-02 | -1.237 | 1.3E-02 |
| WHRN | 0.979 | 1.5E-03 | 0.471 | 1.5E-02 | 0.466 | 1.7E-02 |
| WWC3 | 0.275 | 1.4E-03 | 0.128 | 1.8E-02 | 0.137 | 1.2E-02 |
| XK | -0.674 | 9.6E-03 | 2.073 | 5.6E-03 | -1.728 | 2.3E-02 |
| XRCC3 | 0.578 | 4.3E-03 | 0.324 | 4.2E-03 | 0.254 | 2.4E-02 |
| XYLB | -0.437 | 2.8E-05 | -0.246 | 7.6E-04 | -0.202 | 5.8E-03 |
| YWHAZ | -0.657 | 1.0E-06 | -0.317 | 3.6E-03 | -0.346 | 1.5E-03 |
| ZBED6 | 0.476 | 1.8E-03 | 0.226 | 3.6E-03 | 0.237 | 2.8E-03 |
| ZBTB16 | 0.555 | 1.3E-03 | 0.272 | 3.9E-02 | 0.269 | 4.0E-02 |
| ZC2HC1A | -0.441 | 5.9E-03 | -0.244 | 5.6E-03 | -0.212 | 1.5E-02 |
| ZC3H11A | 0.476 | 1.8E-03 | 0.226 | 3.6E-03 | 0.237 | 2.8E-03 |
| ZCCHC24 | 1.011 | 6.2E-06 | 0.557 | 2.0E-04 | 0.447 | 2.3E-03 |
| ZGPAT | 0.498 | 3.6E-03 | 0.216 | 1.9E-02 | 0.27 | 3.3E-03 |
| ZMIZ1-AS1 | -0.668 | 4.9E-05 | -0.291 | 5.0E-03 | -0.401 | 1.6E-04 |
| ZMYND8 | 0.514 | 6.8E-05 | 0.267 | 3.1E-03 | 0.238 | 9.7E-03 |
| ZNF10 | -0.594 | 8.0E-04 | -0.374 | 3.7E-04 | -0.234 | 2.6E-02 |
| ZNF184 | -0.791 | 1.8E-04 | -0.397 | 7.1E-03 | -0.399 | 6.6E-03 |
| ZNF25 | -0.338 | 9.1E-03 | -0.168 | 4.6E-02 | -0.181 | 3.2E-02 |
| ZNF345 | 0.383 | 9.0E-04 | 0.197 | 1.3E-03 | 0.167 | 9.5E-03 |
| ZNF347 | 0.501 | 1.1E-03 | 0.284 | 1.0E-03 | 0.202 | 1.7E-02 |
| ZNF441 | -0.831 | 3.8E-05 | -0.365 | 2.0E-02 | -0.45 | 4.9E-03 |
| ZNF542P | -0.374 | 2.0E-02 | -0.196 | 3.9E-02 | -0.188 | 4.7E-02 |
| ZNF561-AS1 | 0.357 | 3.4E-03 | 0.17 | 1.6E-02 | 0.177 | 1.1E-02 |
| ZNF592 | 0.268 | 1.2E-02 | 0.137 | 1.9E-02 | 0.127 | 3.2E-02 |
| ZNF652 | 0.929 | 7.3E-04 | 0.442 | 9.5E-03 | 0.486 | 5.1E-03 |
| ZNF672 | 0.406 | 1.4E-02 | 0.193 | 3.2E-02 | 0.204 | 2.1E-02 |
| ZNF787 | 0.464 | 7.5E-03 | 0.231 | 1.3E-02 | 0.225 | 1.9E-02 |
| ZNF860 | -1.53 | 2.4E-03 | -0.918 | 3.6E-03 | -0.642 | 4.0E-02 |
| ZRANB3 | -0.257 | 3.4E-03 | -0.166 | 4.4E-04 | -0.101 | 3.4E-02 |
| ZXDC | 0.316 | 9.7E-03 | 0.151 | 2.3E-02 | 0.155 | 2.0E-02 |

Based on the deseq, and filtered using IPA, genes that were expressed in all COVID-19 and AD datasets with an absolute expression of 0.0001 or greater and p<0.05 are presented in Supplemental Data 2. Expr. Is expression.
