## Supplemental Data 2 for "Molecular and cellular similarities in the brain of SARS-CoV-2 and Alzheimer’s disease individuals"

Supplemental Table 4: Gene regulation within the HF

| | AD+COVID-19 vs. Ctrl | | COVID-19 vs. Ctrl | | AD vs. Ctrl | |
|---|---|---|---|---|---|---|
| Gene | Expr. Log Ratio | Expr. p-value | Expr. Log Ratio | Expr. p-value | Expr. Log Ratio | Expr. p-value |
| ABCA10 | -0.659 | 2.6E-03 | -0.347 | 7.9E-03 | -0.343 | 8.5E-03 |
| ABCC12 | -1.704 | 1.2E-03 | -0.715 | 2.7E-02 | -1.152 | 2.0E-04 |
| ACIN1 | 0.341 | 3.8E-04 | 0.212 | 4.2E-05 | 0.13 | 1.2E-02 |
| ACTR3B | -0.486 | 1.9E-03 | -0.304 | 6.6E-04 | -0.176 | 4.9E-02 |
| ADAMTS2 | 1.192 | 4.3E-04 | 0.539 | 4.1E-02 | 0.669 | 1.1E-02 |
| ADAT3 | 0.757 | 2.3E-03 | 0.4 | 1.1E-02 | 0.349 | 2.5E-02 |
| AEBP1 | 1.094 | 2.5E-03 | 0.574 | 1.4E-02 | 0.511 | 3.0E-02 |
| AGAP6 (inc | 0.538 | 9.9E-03 | 0.296 | 1.7E-02 | 0.241 | 4.9E-02 |
| AHSA2P | 0.554 | 1.4E-03 | 0.332 | 6.6E-04 | 0.225 | 2.1E-02 |
| ALOX15B | 1.954 | 4.9E-03 | 0.94 | 3.5E-02 | 0.905 | 4.5E-02 |
| ANAPC16 | 0.608 | 3.8E-04 | 0.322 | 8.7E-03 | 0.275 | 2.6E-02 |
| ANKRD20A | 1.852 | 3.7E-03 | 1.087 | 1.5E-02 | 0.894 | 4.9E-02 |
| ANKRD20A | -1.324 | 4.5E-03 | -0.857 | 2.7E-02 | -0.955 | 3.0E-03 |
| ANKRD34B | -2.696 | 1.3E-04 | -1.243 | 4.2E-03 | -1.331 | 4.3E-04 |
| APOO | -0.435 | 1.9E-04 | -0.212 | 2.9E-03 | -0.214 | 2.6E-03 |
| ARAP1 | 0.805 | 1.5E-07 | 0.538 | 8.4E-09 | 0.253 | 6.5E-03 |
| ARAP1-AS2 | 0.957 | 7.2E-06 | 0.669 | 2.0E-07 | 0.274 | 3.3E-02 |
| ARHGAP36 | -3.058 | 1.4E-04 | -1.551 | 1.0E-04 | -1.313 | 1.1E-03 |
| ARSG | -1.235 | 1.1E-05 | -0.703 | 5.1E-05 | -0.553 | 1.5E-03 |
| ASAH2B | -1.039 | 2.2E-06 | -0.604 | 3.9E-04 | -0.437 | 8.9E-03 |
| ATG4B | 0.518 | 7.0E-05 | 0.28 | 3.5E-03 | 0.242 | 1.3E-02 |
| ATOH8 | 1.12 | 3.9E-03 | 0.571 | 7.9E-03 | 0.569 | 8.0E-03 |
| ATP6V0E2- | -0.497 | 4.9E-04 | -0.317 | 9.9E-05 | -0.174 | 3.5E-02 |
| AUP1 | 0.549 | 4.9E-04 | 0.303 | 3.5E-03 | 0.237 | 2.0E-02 |
| B3GALNT1 | -0.626 | 1.4E-02 | -0.329 | 9.6E-03 | -0.287 | 2.3E-02 |
| B3GAT1-D1 | -1.023 | 8.0E-04 | -0.501 | 7.2E-03 | -0.608 | 1.1E-03 |
| BAG6 | -3.222 | 8.2E-04 | -2.088 | 3.3E-04 | -1.195 | 4.1E-02 |
| BANP | 0.907 | 5.9E-04 | 0.46 | 1.3E-02 | 0.397 | 3.2E-02 |
| BCLAF3 | -0.332 | 4.0E-04 | -0.16 | 5.2E-03 | -0.166 | 6.2E-03 |
| BLCAP | -0.567 | 3.7E-03 | -0.298 | 3.3E-03 | -0.253 | 1.5E-02 |
| BMP1 | 0.679 | 2.2E-03 | 0.383 | 5.2E-03 | 0.288 | 3.6E-02 |
| BOLA2/BOI | -4.535 | 9.3E-03 | -0.848 | 7.9E-03 | -4.101 | 1.5E-03 |
| BRF1 | 0.41 | 4.1E-05 | 0.211 | 2.8E-04 | 0.201 | 5.4E-04 |
| BUB1B | -1.387 | 8.1E-03 | -0.674 | 2.4E-02 | -0.645 | 3.2E-02 |
| CACNA2D4 | -0.595 | 5.0E-03 | -0.29 | 4.0E-02 | -0.297 | 3.4E-02 |
| CAPN2 | 0.562 | 8.8E-04 | 0.285 | 3.8E-03 | 0.281 | 5.6E-03 |
| CAPZB | 0.318 | 5.8E-04 | 0.13 | 1.3E-02 | 0.191 | 2.5E-04 |
| CARD11 | 1.197 | 4.5E-04 | 0.583 | 3.0E-02 | 0.648 | 1.7E-02 |
| CARHSP1 | 0.595 | 2.5E-05 | 0.351 | 4.7E-05 | 0.241 | 5.2E-03 |
| CASP7 | 1.213 | 4.0E-04 | 0.719 | 1.8E-04 | 0.45 | 2.4E-02 |
| CBS/LOC10 | 2.431 | 1.8E-04 | 0.879 | 2.6E-02 | 1.59 | 6.6E-05 |
| CCDC144N | -0.483 | 1.7E-02 | -0.236 | 2.9E-02 | -0.242 | 2.6E-02 |
| CCDC32 | -0.525 | 5.1E-05 | -0.344 | 4.4E-05 | -0.177 | 3.5E-02 |

| | | | | | | |
|---|---|---|---|---|---|---|
| CCPG1 | -0.423 | 1.8E-03 | -0.263 | 1.1E-04 | -0.16 | 2.1E-02 |
| CCSER2 | -0.257 | 2.2E-04 | -0.14 | 2.4E-03 | -0.112 | 1.9E-02 |
| CD200 | -0.685 | 6.8E-03 | -0.353 | 2.4E-02 | -0.329 | 3.6E-02 |
| CDC42EP2 | 0.894 | 8.3E-04 | 0.445 | 4.0E-02 | 0.453 | 3.7E-02 |
| CDK15 | -1.42 | 1.5E-02 | -0.702 | 2.3E-02 | -0.722 | 1.9E-02 |
| CDK2AP1 | 0.637 | 1.1E-03 | 0.327 | 2.7E-03 | 0.316 | 4.7E-03 |
| CDK6-AS1 | -1.429 | 1.2E-03 | -0.75 | 1.2E-02 | -0.674 | 2.3E-02 |
| CHDH | 0.813 | 4.7E-04 | 0.515 | 4.9E-04 | 0.295 | 4.3E-02 |
| CHEK2 | 1.184 | 1.9E-05 | 0.543 | 2.3E-03 | 0.596 | 8.3E-04 |
| CHKB-CPT1 | 0.671 | 4.9E-04 | 0.37 | 8.4E-03 | 0.314 | 2.7E-02 |
| CHST3 | 0.914 | 4.4E-03 | 0.502 | 3.4E-02 | 0.47 | 4.8E-02 |
| CIP2A | -1.278 | 1.8E-04 | -0.694 | 2.6E-03 | -0.607 | 8.1E-03 |
| CLASRP | 0.582 | 9.7E-04 | 0.382 | 2.9E-05 | 0.197 | 2.5E-02 |
| CLCN2 | 0.535 | 5.3E-03 | 0.241 | 3.4E-02 | 0.294 | 9.8E-03 |
| CLDN15 | 1.169 | 8.6E-08 | 0.797 | 6.4E-07 | 0.361 | 2.3E-02 |
| CLK1 | 0.595 | 1.4E-04 | 0.235 | 2.0E-02 | 0.355 | 4.7E-04 |
| COLGALT1 | 0.743 | 4.8E-05 | 0.375 | 1.6E-03 | 0.366 | 2.5E-03 |
| COPG2 | -0.815 | 1.1E-02 | -0.438 | 1.3E-02 | -0.357 | 4.5E-02 |
| COPS7A | -0.558 | 8.5E-05 | -0.327 | 1.8E-04 | -0.227 | 7.6E-03 |
| COX7B | -0.509 | 1.2E-03 | -0.278 | 8.3E-03 | -0.232 | 2.8E-02 |
| CRYAB | 1.092 | 4.8E-04 | 0.578 | 6.3E-03 | 0.425 | 4.5E-02 |
| CRYBA1 | 0.971 | 2.2E-03 | 0.48 | 3.2E-02 | 0.52 | 2.1E-02 |
| CSRP1 | 0.907 | 3.6E-04 | 0.41 | 1.2E-02 | 0.498 | 2.2E-03 |
| CTBP2 | 0.595 | 5.8E-05 | 0.293 | 7.0E-04 | 0.299 | 5.3E-04 |
| CTF2P | 2.471 | 7.9E-04 | 1.522 | 1.9E-03 | 1.004 | 3.7E-02 |
| CTTN | 0.308 | 1.7E-02 | 0.154 | 3.4E-02 | 0.159 | 2.9E-02 |
| DAZAP1 | 0.577 | 3.8E-05 | 0.339 | 4.0E-05 | 0.235 | 4.5E-03 |
| DDX11L2 | 1.58 | 3.8E-03 | 0.796 | 3.2E-02 | 0.956 | 1.0E-02 |
| DDX23 | 0.416 | 1.7E-02 | 0.199 | 3.6E-02 | 0.224 | 1.8E-02 |
| DEK | 0.557 | 4.5E-04 | 0.259 | 3.4E-03 | 0.294 | 9.0E-04 |
| DGKH | -0.492 | 8.3E-03 | -0.215 | 2.8E-02 | -0.271 | 4.7E-03 |
| DHODH | 0.526 | 4.1E-03 | 0.237 | 2.5E-02 | 0.283 | 7.3E-03 |
| DHX57 | -0.432 | 2.2E-03 | -0.229 | 4.7E-03 | -0.198 | 1.4E-02 |
| DIRAS3 | -1.409 | 2.1E-03 | -0.589 | 1.8E-02 | -0.775 | 1.6E-03 |
| DLD | -0.384 | 1.2E-03 | -0.215 | 1.5E-03 | -0.163 | 1.6E-02 |
| DNAAF4-C( | -0.41 | 2.5E-03 | -0.242 | 1.9E-04 | -0.17 | 9.3E-03 |
| DNAJC5G | -1.504 | 1.9E-03 | -1.009 | 3.0E-04 | -0.563 | 4.0E-02 |
| DNAL4 | -0.694 | 1.8E-03 | -0.368 | 1.0E-02 | -0.32 | 2.6E-02 |
| DOT1L | 0.5 | 4.3E-03 | 0.235 | 4.0E-02 | 0.274 | 1.6E-02 |
| DPF2 | 0.301 | 8.3E-04 | 0.189 | 7.6E-04 | 0.111 | 4.8E-02 |
| DPY19L2P3 | -0.635 | 8.2E-04 | -0.333 | 1.4E-03 | -0.296 | 4.6E-03 |
| DRD3 | -2.669 | 1.1E-02 | -1.256 | 4.6E-02 | -1.295 | 2.7E-02 |
| DSCR9 | -0.877 | 5.0E-03 | -0.419 | 3.0E-02 | -0.435 | 2.2E-02 |
| EFHD1 | 1.023 | 1.7E-03 | 0.521 | 4.1E-02 | 0.544 | 3.4E-02 |
| EGFEM1P | -0.889 | 2.1E-03 | -0.39 | 4.2E-02 | -0.492 | 1.0E-02 |
| EGFL6 | -2.451 | 2.7E-03 | -1.495 | 3.5E-03 | -1.089 | 2.7E-02 |
| EHMT1 | 0.328 | 3.9E-04 | 0.189 | 5.3E-04 | 0.14 | 1.0E-02 |

| | | | | | | |
|---|---|---|---|---|---|---|
| ELF1 | 0.572 | 1.7E-03 | 0.28 | 3.9E-02 | 0.303 | 2.5E-02 |
| ELMOD3 | 0.597 | 3.9E-06 | 0.314 | 6.1E-05 | 0.285 | 2.8E-04 |
| ELOA-AS1 | 0.48 | 2.0E-03 | 0.229 | 1.3E-02 | 0.248 | 6.0E-03 |
| EMBP1 | -0.904 | 5.4E-03 | -0.416 | 3.5E-02 | -0.415 | 3.7E-02 |
| EMP3 | 1.159 | 2.7E-04 | 0.499 | 3.3E-02 | 0.593 | 1.4E-02 |
| EPS8L3 | 2.545 | 5.8E-03 | 1.235 | 3.6E-02 | 1.336 | 2.4E-02 |
| ERCC8 | -0.27 | 1.1E-02 | -0.129 | 4.7E-02 | -0.132 | 4.7E-02 |
| ERI3 | 0.337 | 3.1E-04 | 0.206 | 1.7E-03 | 0.135 | 4.2E-02 |
| FAM133B | 0.538 | 6.5E-04 | 0.206 | 3.9E-02 | 0.331 | 8.4E-04 |
| FAM3D-AS | 1.176 | 8.1E-04 | 0.703 | 2.1E-03 | 0.485 | 3.3E-02 |
| FAM53B | 0.647 | 2.3E-05 | 0.337 | 3.0E-03 | 0.314 | 5.8E-03 |
| FBXO2 | 0.977 | 9.6E-05 | 0.615 | 2.3E-04 | 0.367 | 3.9E-02 |
| FCGR2A | 0.994 | 1.1E-03 | 0.457 | 2.2E-02 | 0.587 | 3.9E-03 |
| FCHO1 | 0.705 | 5.1E-05 | 0.373 | 1.5E-02 | 0.341 | 2.7E-02 |
| FLII | 0.572 | 1.2E-04 | 0.334 | 4.0E-05 | 0.238 | 3.5E-03 |
| FREM3 | -1.182 | 5.4E-04 | -0.607 | 2.3E-03 | -0.619 | 2.0E-03 |
| FSCN2 | 1.526 | 8.4E-05 | 0.952 | 2.8E-04 | 0.58 | 2.7E-02 |
| GAD2 | -1.586 | 1.1E-03 | -0.828 | 1.1E-02 | -0.759 | 1.7E-02 |
| GGT4P | -2.405 | 2.1E-04 | -1.528 | 3.9E-03 | -1.214 | 1.8E-02 |
| GNA13 | 0.638 | 3.5E-04 | 0.304 | 1.5E-03 | 0.324 | 8.8E-04 |
| GNG4 | -1.028 | 1.2E-04 | -0.487 | 1.3E-02 | -0.587 | 1.7E-03 |
| GNRH1 | 0.947 | 2.1E-03 | 0.464 | 4.7E-02 | 0.478 | 4.8E-02 |
| GOLGA8A/ | -0.783 | 2.4E-02 | -0.386 | 4.1E-02 | -0.46 | 5.0E-02 |
| GPR101 | -2.715 | 4.8E-04 | -1.387 | 2.5E-03 | -1.277 | 2.7E-03 |
| GPR39 | -0.875 | 1.8E-04 | -0.473 | 1.3E-03 | -0.384 | 1.0E-02 |
| GPR4 | 1.105 | 2.3E-03 | 0.592 | 1.9E-02 | 0.505 | 4.7E-02 |
| GPRC5B | 0.714 | 9.4E-05 | 0.409 | 4.4E-03 | 0.305 | 3.4E-02 |
| GSTA4 | -0.492 | 7.4E-03 | -0.292 | 8.9E-04 | -0.202 | 2.5E-02 |
| GTF2IP7 | -1.517 | 3.3E-04 | -0.722 | 3.5E-02 | -0.753 | 2.7E-02 |
| HCLS1 | 1.324 | 3.1E-05 | 0.763 | 2.5E-04 | 0.532 | 1.1E-02 |
| HDAC4 | 0.509 | 5.1E-06 | 0.311 | 1.0E-07 | 0.199 | 7.0E-04 |
| HDGF | 0.509 | 1.7E-03 | 0.28 | 4.0E-04 | 0.23 | 4.3E-03 |
| HEXD | 0.875 | 1.2E-07 | 0.519 | 1.5E-05 | 0.368 | 2.5E-03 |
| HMBOX1 | 0.436 | 8.1E-04 | 0.224 | 2.1E-03 | 0.215 | 3.4E-03 |
| HMG20B | 0.733 | 7.8E-03 | 0.358 | 1.5E-02 | 0.368 | 1.2E-02 |
| HNRNPA2B | 0.34 | 8.5E-04 | 0.124 | 3.4E-02 | 0.215 | 2.4E-04 |
| HP1BP3 | 0.392 | 3.8E-03 | 0.202 | 3.3E-02 | 0.19 | 4.4E-02 |
| HS6ST2 | -1.06 | 8.2E-03 | -0.487 | 4.8E-02 | -0.568 | 1.8E-02 |
| ICA1 | -0.861 | 5.8E-03 | -0.428 | 7.4E-03 | -0.477 | 2.9E-03 |
| IKBKB | 0.47 | 6.0E-04 | 0.325 | 1.2E-05 | 0.146 | 4.8E-02 |
| IL1RL2 | -0.998 | 9.9E-04 | -0.45 | 2.1E-02 | -0.562 | 4.5E-03 |
| INSR | 0.408 | 6.8E-04 | 0.251 | 6.1E-04 | 0.162 | 2.6E-02 |
| IP6K2 | 0.348 | 5.6E-03 | 0.175 | 2.9E-02 | 0.177 | 2.8E-02 |
| IRS2 | 0.644 | 4.1E-03 | 0.299 | 4.1E-02 | 0.343 | 1.9E-02 |
| ITGA11 | 1.186 | 9.3E-05 | 0.749 | 1.0E-04 | 0.461 | 2.0E-02 |
| ITGB8-AS1 | 0.865 | 1.4E-03 | 0.532 | 1.2E-03 | 0.359 | 2.9E-02 |
| ITPKB | 0.855 | 8.8E-04 | 0.375 | 4.5E-02 | 0.505 | 6.9E-03 |

| | | | | | | |
|---|---|---|---|---|---|---|
| IWS1 | 0.446 | 6.7E-05 | 0.182 | 3.7E-02 | 0.266 | 2.6E-03 |
| JAKMIP3 | 0.791 | 5.4E-05 | 0.389 | 8.4E-03 | 0.403 | 6.3E-03 |
| JPT1 | -0.503 | 4.9E-04 | -0.281 | 9.9E-04 | -0.216 | 1.1E-02 |
| KCNE4 | 1.205 | 2.9E-04 | 0.59 | 8.1E-03 | 0.604 | 7.5E-03 |
| KLC1 | 0.194 | 8.9E-03 | 0.102 | 2.5E-02 | 0.095 | 3.7E-02 |
| KLF15 | 1.156 | 2.0E-04 | 0.784 | 1.6E-05 | 0.385 | 3.3E-02 |
| KLHDC10 | -0.396 | 8.7E-05 | -0.26 | 6.3E-06 | -0.129 | 2.9E-02 |
| KLHL21 | 0.705 | 1.4E-03 | 0.396 | 4.7E-03 | 0.299 | 3.4E-02 |
| KLHL42 | -0.423 | 2.2E-03 | -0.264 | 2.7E-04 | -0.155 | 3.2E-02 |
| LANCL2 | -0.476 | 1.8E-03 | -0.217 | 2.5E-02 | -0.26 | 7.9E-03 |
| LBX2 | 1.477 | 1.6E-03 | 0.741 | 1.9E-02 | 0.694 | 2.7E-02 |
| LCLAT1 | -0.489 | 1.6E-04 | -0.308 | 5.3E-04 | -0.181 | 4.3E-02 |
| LDAH | -0.463 | 1.6E-02 | -0.232 | 2.7E-02 | -0.231 | 3.0E-02 |
| LENG8 | -3.922 | 6.4E-05 | -2.389 | 1.8E-04 | -2.345 | 3.0E-04 |
| LINC00390 | -1.426 | 1.5E-03 | -0.62 | 2.6E-02 | -0.835 | 2.0E-03 |
| LINC00484 | 1.145 | 2.6E-02 | 0.48 | 1.5E-02 | 0.737 | 2.7E-02 |
| LINC00501 | -1.537 | 5.5E-03 | -0.62 | 3.8E-02 | -0.999 | 1.2E-03 |
| LINC00842 | -1.61 | 7.6E-04 | -0.791 | 2.1E-02 | -0.816 | 1.6E-02 |
| LINC00862 | 1.033 | 6.9E-04 | 0.538 | 1.3E-02 | 0.438 | 4.3E-02 |
| LINC00921 | 1.149 | 2.0E-03 | 0.475 | 4.2E-02 | 0.568 | 1.4E-02 |
| LINC00923 | -1.153 | 3.3E-03 | -0.596 | 1.3E-02 | -0.507 | 3.7E-02 |
| LINC01013 | -1.119 | 1.3E-03 | -0.51 | 3.2E-02 | -0.64 | 7.6E-03 |
| LINC01090 | -1.252 | 1.1E-03 | -0.765 | 2.5E-03 | -0.498 | 4.8E-02 |
| LINC01134 | 1.735 | 2.9E-03 | 0.993 | 6.1E-03 | 0.767 | 3.4E-02 |
| LINC01168 | -2.347 | 1.1E-04 | -0.948 | 1.3E-02 | -1.374 | 1.7E-04 |
| LINC01277 | -0.984 | 1.8E-03 | -0.538 | 7.7E-03 | -0.47 | 2.0E-02 |
| LINC01310 | -1.128 | 2.3E-03 | -0.563 | 2.0E-02 | -0.646 | 7.5E-03 |
| LINC01362 | -0.862 | 7.6E-03 | -0.432 | 3.2E-02 | -0.432 | 3.0E-02 |
| LINC01440 | -2.492 | 1.3E-05 | -1.229 | 4.7E-03 | -1.247 | 2.5E-03 |
| LINC01537 | -1.536 | 1.3E-03 | -0.538 | 4.4E-02 | -1.061 | 1.3E-04 |
| LINC01730 | 1.862 | 2.6E-03 | 0.786 | 3.6E-02 | 1.02 | 6.7E-03 |
| LINC01736 | 2.078 | 1.4E-03 | 1.16 | 1.2E-02 | 0.956 | 3.9E-02 |
| LINC01807 | -1.15 | 1.4E-03 | -0.635 | 1.1E-02 | -0.492 | 4.7E-02 |
| LINC01878 | 1.301 | 2.8E-03 | 0.566 | 4.1E-02 | 0.721 | 9.4E-03 |
| LINC02008 | -1.241 | 9.9E-05 | -0.514 | 4.4E-02 | -0.763 | 2.3E-03 |
| LINC02211 | -5.003 | 1.8E-04 | -1.979 | 1.0E-02 | -3.177 | 3.6E-05 |
| LINC02320 | -1.546 | 1.7E-04 | -0.728 | 2.0E-03 | -0.868 | 2.0E-04 |
| LINC02397 | 3.053 | 5.3E-05 | 1.997 | 2.6E-03 | 1.506 | 2.3E-02 |
| LINC02752 | -1.92 | 3.4E-07 | -0.915 | 4.2E-03 | -1.126 | 2.6E-04 |
| LINC02756 | -2.046 | 9.3E-03 | -1.028 | 2.8E-02 | -1.097 | 1.7E-02 |
| LINC02774 | -1.128 | 3.0E-03 | -0.462 | 2.5E-02 | -0.697 | 8.0E-04 |
| LNCAROD | 1.749 | 6.1E-05 | 1.034 | 2.0E-03 | 0.773 | 2.1E-02 |
| LYRM9 | -1.17 | 3.2E-05 | -0.632 | 5.7E-04 | -0.566 | 2.1E-03 |
| MAD1L1 | 0.764 | 1.6E-04 | 0.502 | 2.2E-06 | 0.249 | 2.0E-02 |
| MAGED1 | -0.492 | 2.7E-04 | -0.333 | 1.4E-06 | -0.157 | 4.5E-02 |
| MAP3K13 | -0.44 | 2.7E-04 | -0.267 | 1.2E-03 | -0.173 | 3.5E-02 |
| MAP4K3-D' | -0.603 | 2.0E-02 | -0.346 | 7.5E-03 | -0.265 | 4.1E-02 |

| | | | | | | |
|---|---|---|---|---|---|---|
| MCMDC2 | 1.328 | 5.5E-04 | 0.524 | 4.7E-02 | 0.739 | 5.1E-03 |
| MEAK7 | -0.576 | 7.1E-04 | -0.312 | 4.6E-03 | -0.261 | 1.7E-02 |
| MEG9 | -1.031 | 9.5E-03 | -0.514 | 1.9E-02 | -0.549 | 1.4E-02 |
| MEP1A | 2.708 | 3.2E-03 | 1.196 | 4.3E-02 | 1.748 | 3.4E-03 |
| MEST | -0.801 | 9.2E-03 | -0.4 | 2.2E-02 | -0.374 | 3.3E-02 |
| METAP1D | -0.62 | 2.6E-03 | -0.335 | 1.3E-02 | -0.28 | 3.8E-02 |
| MIA | 2.044 | 3.9E-03 | 1.13 | 1.2E-02 | 1.046 | 2.0E-02 |
| MIA-RAB4E | 0.453 | 6.8E-03 | 0.214 | 4.5E-02 | 0.226 | 3.2E-02 |
| MID1IP1 | 0.94 | 2.4E-05 | 0.477 | 1.9E-03 | 0.463 | 2.5E-03 |
| MOB1A | 0.377 | 1.9E-03 | 0.19 | 1.9E-02 | 0.183 | 2.4E-02 |
| MS4A14 | 1.595 | 1.4E-04 | 0.962 | 5.9E-04 | 0.622 | 2.7E-02 |
| MS4A6E | 1.299 | 5.7E-04 | 0.764 | 3.4E-03 | 0.581 | 2.7E-02 |
| MS4A7 | 1.27 | 9.7E-04 | 0.776 | 3.0E-03 | 0.535 | 4.1E-02 |
| MS4A8 | -2.386 | 5.4E-05 | -1.081 | 2.8E-03 | -1.153 | 1.4E-03 |
| MT1E | 1.264 | 1.3E-06 | 0.88 | 1.2E-06 | 0.387 | 3.4E-02 |
| MT1F | 1.567 | 5.2E-07 | 1.103 | 6.2E-07 | 0.473 | 3.2E-02 |
| MT1G | 1.887 | 4.4E-07 | 1.301 | 1.7E-07 | 0.574 | 2.6E-02 |
| MTCO3P12 | -1.252 | 5.1E-03 | -0.708 | 3.0E-02 | -0.679 | 3.3E-02 |
| MTMR14 | 0.35 | 1.1E-02 | 0.203 | 2.8E-03 | 0.146 | 3.4E-02 |
| NBPF10 (in | 0.62 | 1.5E-03 | 0.343 | 3.4E-02 | 0.278 | 4.9E-02 |
| NCR3LG1 | -0.706 | 7.1E-04 | -0.403 | 2.3E-03 | -0.315 | 1.7E-02 |
| NDUFA5 | -0.718 | 5.7E-05 | -0.5 | 7.4E-07 | -0.209 | 4.4E-02 |
| NDUFAF6 | 0.459 | 1.0E-03 | 0.262 | 4.9E-03 | 0.195 | 3.7E-02 |
| NEAT1 | 1.378 | 3.9E-05 | 0.464 | 2.6E-02 | 0.9 | 1.7E-05 |
| NKX6-2 | 1.269 | 5.6E-08 | 0.848 | 7.0E-05 | 0.441 | 3.9E-02 |
| NLRP3P1 | -1.374 | 5.6E-03 | -0.626 | 4.9E-02 | -0.82 | 1.0E-02 |
| NMD3 | -0.64 | 5.9E-04 | -0.244 | 2.6E-02 | -0.401 | 2.2E-04 |
| NRIR | -1.058 | 8.4E-04 | -0.592 | 1.5E-02 | -0.505 | 3.6E-02 |
| NRSN2 | -0.619 | 8.0E-04 | -0.264 | 2.7E-02 | -0.358 | 2.8E-03 |
| NTNG1 | -0.793 | 4.9E-06 | -0.478 | 1.7E-04 | -0.318 | 1.1E-02 |
| NUP188 | 0.329 | 3.0E-03 | 0.175 | 7.3E-03 | 0.151 | 2.5E-02 |
| NUP42 | -0.668 | 6.6E-04 | -0.287 | 1.3E-02 | -0.379 | 9.4E-04 |
| OLMALINC | 0.948 | 5.4E-04 | 0.38 | 2.5E-02 | 0.565 | 1.5E-03 |
| OR2AG2 | -1.313 | 6.6E-03 | -0.702 | 2.2E-02 | -0.748 | 1.5E-02 |
| P2RX7 | 0.566 | 1.1E-04 | 0.32 | 4.4E-03 | 0.25 | 2.6E-02 |
| PACS2 | 0.518 | 1.8E-03 | 0.296 | 1.9E-03 | 0.228 | 1.7E-02 |
| PALLD | 0.744 | 6.5E-03 | 0.4 | 1.1E-02 | 0.322 | 4.3E-02 |
| PARVG | 1.36 | 2.0E-07 | 0.884 | 1.1E-07 | 0.487 | 5.2E-03 |
| PCDH1 | -0.576 | 1.1E-03 | -0.264 | 8.2E-03 | -0.306 | 2.1E-03 |
| PCID2 | 0.471 | 4.5E-03 | 0.238 | 1.9E-02 | 0.236 | 2.1E-02 |
| PCSK1 | -1.09 | 4.8E-05 | -0.734 | 1.1E-04 | -0.374 | 4.9E-02 |
| PCYOX1L | -1.098 | 1.7E-04 | -0.559 | 1.7E-03 | -0.59 | 9.1E-04 |
| PENK | -4.081 | 4.2E-04 | -1.81 | 1.6E-02 | -2.247 | 1.6E-03 |
| PHF19 | 0.968 | 1.6E-06 | 0.543 | 1.3E-03 | 0.437 | 9.7E-03 |
| PIAS4 | 0.685 | 4.5E-08 | 0.414 | 1.2E-07 | 0.276 | 4.3E-04 |
| PID1 | -0.596 | 1.5E-03 | -0.266 | 2.8E-02 | -0.33 | 6.0E-03 |
| PIPOX | -0.775 | 5.6E-06 | -0.243 | 2.2E-02 | -0.534 | 2.4E-07 |

| | | | | | | |
|---|---|---|---|---|---|---|
| PKN2-AS1 | -0.695 | 7.0E-03 | -0.322 | 2.6E-02 | -0.367 | 1.1E-02 |
| PLIN3 | 0.985 | 2.3E-03 | 0.475 | 3.6E-02 | 0.548 | 1.5E-02 |
| PLOD3 | 1.045 | 5.7E-05 | 0.536 | 3.1E-04 | 0.432 | 3.8E-03 |
| PLXNB1 | 0.841 | 1.7E-05 | 0.536 | 5.3E-05 | 0.314 | 1.7E-02 |
| PMS2P4 | -0.78 | 6.4E-04 | -0.311 | 2.3E-02 | -0.462 | 7.0E-04 |
| POLD1 | 0.847 | 3.0E-04 | 0.516 | 1.1E-04 | 0.313 | 1.6E-02 |
| POLR2H | 0.45 | 4.6E-03 | 0.272 | 2.8E-03 | 0.189 | 4.0E-02 |
| PPP1R2 | -0.597 | 1.7E-05 | -0.416 | 4.9E-06 | -0.18 | 4.8E-02 |
| PPP4R3A | 0.245 | 2.1E-03 | 0.115 | 3.8E-02 | 0.133 | 1.7E-02 |
| PRDM16 | 0.611 | 2.1E-02 | 0.288 | 4.9E-02 | 0.332 | 2.5E-02 |
| PRDX6 | 0.482 | 9.7E-04 | 0.255 | 6.9E-03 | 0.226 | 1.8E-02 |
| PREX1 | 0.796 | 5.8E-05 | 0.489 | 1.4E-03 | 0.315 | 3.9E-02 |
| PRKX | 0.993 | 6.1E-04 | 0.44 | 2.4E-02 | 0.573 | 3.8E-03 |
| QDPR | 0.937 | 1.9E-04 | 0.445 | 1.4E-02 | 0.464 | 1.1E-02 |
| RAB29 | 0.717 | 3.8E-03 | 0.366 | 6.4E-03 | 0.335 | 1.3E-02 |
| RAB41 | 1.092 | 3.6E-03 | 0.57 | 8.2E-03 | 0.442 | 4.2E-02 |
| RAB9A | 0.494 | 5.5E-03 | 0.237 | 2.5E-02 | 0.261 | 1.4E-02 |
| RAF1 | 0.3 | 3.6E-03 | 0.123 | 3.5E-02 | 0.177 | 1.5E-03 |
| RALGAPA2 | 0.382 | 3.1E-04 | 0.185 | 9.2E-03 | 0.201 | 4.7E-03 |
| RANBP3 | 0.5 | 2.9E-04 | 0.311 | 9.6E-05 | 0.189 | 2.0E-02 |
| RASAL3 | 1.527 | 4.0E-05 | 0.998 | 3.3E-05 | 0.526 | 2.9E-02 |
| RBFOX2 | -0.406 | 2.6E-03 | -0.25 | 9.2E-04 | -0.151 | 4.6E-02 |
| RELA | 0.803 | 2.0E-04 | 0.475 | 4.8E-04 | 0.312 | 2.1E-02 |
| RENBP | 1.087 | 4.0E-06 | 0.517 | 2.9E-04 | 0.566 | 1.0E-04 |
| RGCC | 0.891 | 7.4E-04 | 0.501 | 1.2E-02 | 0.43 | 3.0E-02 |
| RGS17 | -0.604 | 3.5E-03 | -0.317 | 3.3E-03 | -0.299 | 5.4E-03 |
| RHOQ | 0.628 | 8.2E-04 | 0.238 | 4.8E-02 | 0.387 | 1.4E-03 |
| RNF169 | 0.4 | 3.8E-04 | 0.222 | 8.2E-03 | 0.177 | 3.4E-02 |
| RNF216 | 0.25 | 1.5E-02 | 0.107 | 3.4E-02 | 0.141 | 5.4E-03 |
| RNFT2 | -0.519 | 7.2E-03 | -0.224 | 4.1E-02 | -0.292 | 8.4E-03 |
| RPH3A | -1.321 | 1.9E-03 | -0.731 | 4.0E-03 | -0.606 | 1.7E-02 |
| RPS6KA1 | 0.781 | 7.6E-04 | 0.431 | 1.8E-02 | 0.358 | 4.8E-02 |
| RPTOR | 0.313 | 3.0E-03 | 0.166 | 5.0E-03 | 0.149 | 1.2E-02 |
| RREB1 | 0.591 | 3.3E-03 | 0.305 | 2.1E-02 | 0.28 | 3.6E-02 |
| RSAD1 | 0.598 | 1.8E-03 | 0.321 | 1.3E-02 | 0.271 | 3.8E-02 |
| RTF1 | -0.448 | 5.4E-05 | -0.285 | 7.2E-05 | -0.155 | 3.3E-02 |
| RUNX1 | 1.073 | 9.0E-08 | 0.615 | 2.2E-06 | 0.47 | 3.1E-04 |
| SAFB | 0.335 | 3.2E-05 | 0.137 | 8.5E-03 | 0.2 | 1.3E-04 |
| SAFB2 | 0.518 | 1.3E-03 | 0.281 | 7.3E-03 | 0.242 | 2.2E-02 |
| SAP30 | 0.608 | 9.5E-04 | 0.253 | 2.6E-02 | 0.37 | 1.4E-03 |
| SCN7A | -0.837 | 1.8E-02 | -0.488 | 1.2E-02 | -0.369 | 4.7E-02 |
| SEC22A | -0.458 | 5.2E-05 | -0.285 | 2.2E-06 | -0.168 | 8.6E-03 |
| SEC24B-AS | -0.783 | 9.8E-03 | -0.399 | 2.3E-02 | -0.416 | 1.7E-02 |
| SEPTIN8 | 0.545 | 3.6E-04 | 0.33 | 2.0E-03 | 0.23 | 3.1E-02 |
| SEZ6 | -0.933 | 1.4E-04 | -0.35 | 2.0E-02 | -0.585 | 8.7E-05 |
| SH2D6 | 1.592 | 1.2E-04 | 0.555 | 2.6E-02 | 1.052 | 3.7E-05 |
| SHKBP1 | 0.805 | 1.6E-03 | 0.47 | 2.8E-03 | 0.329 | 3.7E-02 |

| | | | | | | |
|---|---|---|---|---|---|---|
| SHMT1 | 0.86 | 1.7E-04 | 0.377 | 9.8E-03 | 0.486 | 8.8E-04 |
| SIRPB2 | 1.307 | 6.6E-05 | 0.714 | 5.4E-03 | 0.595 | 1.9E-02 |
| SLC22A23 | 0.58 | 1.7E-04 | 0.27 | 6.4E-03 | 0.326 | 1.5E-03 |
| SLC26A7 | 1.089 | 2.1E-03 | 0.441 | 4.9E-02 | 0.606 | 6.7E-03 |
| SLC2A5 | 1.212 | 1.4E-03 | 0.607 | 1.8E-02 | 0.711 | 4.7E-03 |
| SLC39A11 | 1.014 | 3.7E-08 | 0.359 | 3.3E-02 | 0.667 | 8.5E-05 |
| SLC7A2 | 1.237 | 1.0E-05 | 0.624 | 1.2E-02 | 0.653 | 9.3E-03 |
| SLC8A3 | -1.045 | 2.0E-03 | -0.413 | 3.4E-02 | -0.651 | 7.0E-04 |
| SMAD1-AS: | 1.226 | 1.5E-03 | 0.61 | 1.1E-02 | 0.587 | 1.3E-02 |
| SMG5 | 0.39 | 1.4E-02 | 0.235 | 7.9E-04 | 0.161 | 2.3E-02 |
| SMG9 | 0.571 | 6.6E-04 | 0.315 | 8.5E-03 | 0.254 | 3.5E-02 |
| SMIM1 | 1.859 | 3.8E-04 | 0.835 | 8.0E-03 | 0.89 | 5.6E-03 |
| SMPD4 | 0.37 | 5.0E-05 | 0.23 | 8.2E-05 | 0.141 | 1.4E-02 |
| SNHG10 | 0.828 | 2.0E-04 | 0.444 | 4.1E-03 | 0.383 | 1.5E-02 |
| SNRK | -0.373 | 3.4E-05 | -0.137 | 6.2E-03 | -0.232 | 2.4E-06 |
| SOWAHB | -1.091 | 1.7E-02 | -0.601 | 2.7E-02 | -0.566 | 3.7E-02 |
| SPACA6 | 0.622 | 4.9E-03 | 0.292 | 4.0E-02 | 0.327 | 2.1E-02 |
| SPACA6-AS | 0.768 | 2.2E-03 | 0.383 | 2.0E-02 | 0.397 | 1.6E-02 |
| SPR | 1.247 | 2.3E-04 | 0.701 | 6.4E-03 | 0.534 | 3.8E-02 |
| SPTSSB | -1.401 | 6.3E-04 | -0.782 | 5.0E-04 | -0.675 | 2.8E-03 |
| SRGAP1 | 0.478 | 8.3E-04 | 0.275 | 9.7E-04 | 0.204 | 1.4E-02 |
| SST | -2.468 | 6.0E-06 | -1.47 | 7.4E-04 | -0.951 | 3.0E-02 |
| STARD10 | 0.739 | 9.0E-04 | 0.443 | 7.5E-04 | 0.293 | 2.7E-02 |
| STARD3 | 0.596 | 1.4E-06 | 0.422 | 1.5E-06 | 0.171 | 5.0E-02 |
| STARD4-AS | -1.051 | 3.7E-05 | -0.6 | 1.5E-04 | -0.474 | 2.4E-03 |
| STPG4 | 0.61 | 2.9E-03 | 0.337 | 9.8E-03 | 0.271 | 3.8E-02 |
| STS | -0.582 | 2.3E-02 | -0.313 | 6.6E-03 | -0.284 | 1.3E-02 |
| STUM | -0.905 | 1.6E-03 | -0.362 | 2.5E-02 | -0.536 | 9.2E-04 |
| SULF2 | -0.622 | 5.3E-03 | -0.323 | 2.0E-02 | -0.313 | 2.4E-02 |
| SUN2 | 0.67 | 5.3E-05 | 0.355 | 2.5E-02 | 0.337 | 3.5E-02 |
| SUN3 | -1.97 | 2.8E-04 | -0.957 | 9.8E-03 | -1.146 | 2.3E-03 |
| SV2C | -1.794 | 1.3E-03 | -0.861 | 8.0E-03 | -1.026 | 7.4E-04 |
| SWI5 | 0.373 | 9.3E-03 | 0.183 | 5.0E-03 | 0.175 | 7.7E-03 |
| SYNM | 0.703 | 3.0E-02 | 0.358 | 4.9E-02 | 0.376 | 3.8E-02 |
| SYTL4 | 1.457 | 1.4E-04 | 0.714 | 8.9E-03 | 0.754 | 5.8E-03 |
| TAF1C | 0.746 | 9.6E-08 | 0.485 | 2.8E-06 | 0.274 | 9.6E-03 |
| TARBP1 | -0.769 | 1.8E-04 | -0.395 | 6.4E-05 | -0.38 | 1.5E-04 |
| TASP1 | -0.543 | 2.7E-04 | -0.329 | 1.2E-03 | -0.213 | 3.5E-02 |
| TBCC | -0.626 | 2.9E-03 | -0.295 | 3.7E-02 | -0.314 | 2.9E-02 |
| TBX19 | 0.7 | 2.8E-03 | 0.334 | 4.4E-02 | 0.367 | 2.4E-02 |
| TBX6 | 1.48 | 1.6E-03 | 0.72 | 5.9E-03 | 0.681 | 9.0E-03 |
| TDG | -0.52 | 1.2E-04 | -0.25 | 1.1E-02 | -0.262 | 7.8E-03 |
| TECTA | -0.744 | 9.4E-07 | -0.48 | 1.6E-05 | -0.262 | 1.9E-02 |
| TEMN3-AS: | -2.589 | 5.6E-04 | -1.269 | 2.0E-03 | -1.143 | 7.2E-03 |
| TENM3-AS: | -1.305 | 3.3E-04 | -0.666 | 2.2E-03 | -0.592 | 6.0E-03 |
| TGFB3 | 0.697 | 2.5E-02 | 0.408 | 1.2E-02 | 0.324 | 4.5E-02 |
| TLR5 | 1.043 | 1.0E-03 | 0.544 | 7.9E-04 | 0.518 | 1.2E-03 |

| | | | | | | |
|---|---|---|---|---|---|---|
| TMC6 | 1.059 | 1.8E-05 | 0.562 | 1.0E-02 | 0.515 | 1.9E-02 |
| TMCC2 | 0.61 | 3.5E-03 | 0.3 | 3.3E-02 | 0.306 | 3.1E-02 |
| TMEM53 | 0.504 | 7.4E-05 | 0.252 | 2.5E-02 | 0.259 | 2.3E-02 |
| TPD52L1 | 0.693 | 3.9E-03 | 0.325 | 3.7E-02 | 0.365 | 2.1E-02 |
| TRIM54 | -1.546 | 2.1E-04 | -0.851 | 8.7E-04 | -0.769 | 2.3E-03 |
| TRIM9 | -0.506 | 6.2E-03 | -0.272 | 9.2E-03 | -0.231 | 2.5E-02 |
| TRPC1 | -0.379 | 3.1E-04 | -0.209 | 1.0E-03 | -0.165 | 1.1E-02 |
| TTC31 | 0.42 | 1.7E-02 | 0.203 | 4.5E-02 | 0.213 | 3.5E-02 |
| TUBB1 | 1.079 | 1.9E-03 | 0.475 | 2.8E-02 | 0.659 | 2.1E-03 |
| TUBB3 | -0.951 | 5.1E-04 | -0.509 | 1.0E-02 | -0.414 | 3.8E-02 |
| UBE2D3-AS | 0.738 | 1.6E-03 | 0.408 | 7.8E-03 | 0.336 | 2.7E-02 |
| UBLCP1 | -0.793 | 4.3E-08 | -0.533 | 2.4E-10 | -0.249 | 6.1E-03 |
| UBXN2A | 0.536 | 3.2E-04 | 0.251 | 4.4E-03 | 0.283 | 1.3E-03 |
| UCKL1 | 0.701 | 2.3E-05 | 0.466 | 4.1E-07 | 0.234 | 1.0E-02 |
| UHRF1 | 0.89 | 1.3E-04 | 0.508 | 7.6E-03 | 0.39 | 3.7E-02 |
| UNG | 0.558 | 3.5E-03 | 0.292 | 6.1E-03 | 0.257 | 2.1E-02 |
| UROS | -0.511 | 4.5E-03 | -0.244 | 3.3E-02 | -0.272 | 1.5E-02 |
| USF2 | 0.361 | 9.8E-04 | 0.225 | 1.0E-03 | 0.139 | 4.5E-02 |
| VEZT | -0.379 | 2.6E-05 | -0.249 | 2.5E-06 | -0.125 | 2.0E-02 |
| VGF | -2.426 | 1.4E-07 | -1.405 | 2.5E-04 | -1.065 | 4.9E-03 |
| WWC3 | 0.266 | 1.4E-02 | 0.14 | 1.2E-02 | 0.13 | 2.0E-02 |
| XK | 2.164 | 1.2E-02 | 0.983 | 1.8E-02 | 2.971 | 2.0E-02 |
| XYLB | -0.47 | 1.7E-03 | -0.222 | 4.6E-03 | -0.244 | 1.9E-03 |
| ZBTB16 | 0.652 | 3.1E-03 | 0.28 | 3.4E-02 | 0.379 | 3.9E-03 |
| ZBTB7B | 0.781 | 3.5E-04 | 0.428 | 1.5E-02 | 0.356 | 4.5E-02 |
| ZDHHC20-I | 1.471 | 2.4E-03 | 0.661 | 2.8E-02 | 0.754 | 1.2E-02 |
| ZNF10 | -0.722 | 4.6E-04 | -0.429 | 2.1E-04 | -0.293 | 1.1E-02 |
| ZNF524 | 0.785 | 2.2E-03 | 0.479 | 1.9E-03 | 0.304 | 4.8E-02 |
| ZNF682 | -0.888 | 7.0E-06 | -0.53 | 2.9E-04 | -0.33 | 2.5E-02 |
| ZNF711 | -0.611 | 3.5E-03 | -0.326 | 8.4E-03 | -0.289 | 1.8E-02 |
| ZNF750 | 0.848 | 3.8E-03 | 0.509 | 2.9E-03 | 0.338 | 4.8E-02 |
| ZNF877P | 2.361 | 1.4E-02 | 1.407 | 2.2E-02 | 1.23 | 4.1E-02 |
| ZPR1P1 | 1.179 | 1.0E-03 | 0.592 | 3.4E-02 | 0.62 | 2.7E-02 |
| ZXDC | 0.338 | 8.3E-04 | 0.165 | 1.5E-02 | 0.175 | 9.6E-03 |

Based on the deseq, and filtered using IPA, genes that were expressed in all COVID-19 and AD datasets with an absolute expression of 0.0001 or greater and $p<0.05$ are presented in Suplemental Data 2. Expr. Is expression.
